# Quantifying extinction potential from invasive alien species

**DOI:** 10.1101/2024.09.01.610685

**Authors:** Martin Philippe-Lesaffre, Ugo Arbieu, Alok Bang, Morelia Camacho-Cervantes, Ross N. Cuthbert, Piero Genovesi, Sabrina Kumschick, Arman Pili, Hanno Seebens, Shengyu Wang, Guillaume Latombe

## Abstract

Biological invasions threaten biodiversity, ecosystem services, human health, and cultural heritage, yet their impacts are often underappreciated, leading to insufficient management efforts and suboptimal conservation results. We argue that the lack of quantitative, continuous metrics of impact of invasive alien species (IAS) contribute this lack of appreciation. To bridge this knowledge-action gap, we propose the Extinction Potential Metric (EPM), a suite of quantitative metrics designed to assess the ecological damage caused by IAS. The EPM score of an IAS is the number of current and future species extinctions attributable to this IAS over the next 50 years under a business-as-usual scenario. EPM includes three variants: EPM-A (absolute EPM), EPM-R (relative EPM, which accounts for other anthropogenic pressures), and EPM-U (EPM for Unique species, adjusted for phylogenetic uniqueness of impacted native species), to capture different dimensions of IAS impacts.

We applied EPM to evaluate the impact of IAS on 2178 amphibians, 920 birds, 865 reptiles, and 473 mammals. Our analyses revealed that the impact of the worst IAS was between 90 and 380 times higher than any IAS with an impact of 1 extinct native species. Importantly, several of the most impactful IAS disproportionately affect evolutionarily unique native species.

The EPM framework offers a standardised approach for measuring ecological impacts of IAS but also other anthropogenic pressures at different spatial, temporal, and taxonomic scales. EPM could also guide the development of standardised indicators for assessing the impacts of other anthropogenic stressors. Ultimately, EPM will pave the way to answer ecological questions important to design better conservation policies, to enhance the management of biological invasions and reach global biodiversity conservation goals.

## Introduction

Biological invasions negatively impact many aspects of the natural and human world, including biodiversity, ecosystem services, human health hazards, and indigenous and cultural practices (IPBES 2023). The number of newly introduced alien species has been increasing since the 1800s (Seebens et al. 2017b), and the number of established alien species is expected to increase by 36% between 2005 and 2050 under a business-as-usual scenario (Seebens et al. 2021). The subset of these species with negative impacts, invasive alien species (IAS) hereafter, have contributed to 60% of known species extinctions, cost hundreds of billions of US dollars each year ($423 billion in 2019), and adversely impact quality of human life in 85% of cases (IPBES 2023). The impacts of biological invasions are multiple and complex, and nonetheless remain underestimated and poorly understood by both the public and policy-makers, hindering public support and participation, management actions, and conservation outcomes (Courchamp et al. 2017).

The *InvaCost* database provides a quantitative metric documenting the economic impacts of IAS quantified as monetary costs in a standardised currency and year (Diagne et al. 2020, 2021), succeeding in facilitating policy awareness (Ahmed et al. 2023). For ecological impacts, multiple metrics have been proposed, primarily based on impact categories (see Bernardo-Madrid et al. 2022 for a comparison of seven impact classification schemes). For example, the Environmental Impact Classification for Alien Taxa scheme (EICAT; Blackburn et al. 2014; Hawkins et al. 2015) categorises the maximum global impact of an IAS into five categories, ranging from impacts on individual fitness to the local extinction of species and irreversible changes in community composition, and has been adopted by the IUCN (IUCN 2020a, b). In addition, rankings of IAS have been established (Bellard et al. 2016b), based on the number of threatened species recorded by the IUCN Red List of Threatened Species framework (IUCN 2024) and the Global Invasive Species Database (Pagad et al. 2015). This information has been used at the island level to provide recommendations for eradication (Bellard et al. 2017). Metrics have also been proposed to score and compare different native species impacted by IAS based on their probability of extinction, and their functional distinctiveness and uniqueness, providing important information for their conservation (Marino et al. 2024).

However, to our knowledge, there is no fully quantitative and continuous metric that would allow for comparison of the ecological impacts different IAS have across taxa and scales. Categorical data provide a straightforward tool for managers to assess the threat caused by IAS. EICAT provides managers clear and unambiguous information on the impact of IAS and helps with decision-making and with the prioritisation of IAS management based on the impact categories. However, categorical frameworks are complex to incorporate in statistical analyses that would help us, for example, to quantify the link between ecological impacts and invasion drivers or alien species traits (see Table 1 for a non-exhaustive list of examples of ecological questions and analyses that require a quantitative metric of impact). Existing approaches have for example considered a crude classification into low and high impacts (Evans et al. 2018), or used global ranges of IAS as a proxy for impacts (Byers et al. 2015). Analyses using quantitative impacts would provide a better understanding of how impact varies in space and time and allow for targeted management actions. Moreover, they would allow for more precise profiling of impactful invaders with the aim to forecast future damaging IAS. Since managing biological invasions is among the most effective conservation actions (Langhammer et al. 2024), robust and comparable quantitative metrics and analyses unveiling impact mechanisms would contribute to the prioritisation of actions explicitly required by Target 6 of the Kunming-Montreal Global Biodiversity Framework (GBF), which aims to “eliminate, minimise, reduce and or mitigate the impacts of IAS on biodiversity and ecosystem services”. They would also help achieve Target 4 to “halt species extinction, protect genetic diversity, and manage human-wildlife conflicts” (CBD 2022), and would underpin indicators appropriate for risk identification and prioritisation efforts to meet global biodiversity targets (Butchart et al. 2005; Vicente et al. 2022). Finally, such standardised metrics capable of capturing the severity of ecological impacts of IAS would improve communication between scientists, stakeholders, and policymakers.

**Table 1.**
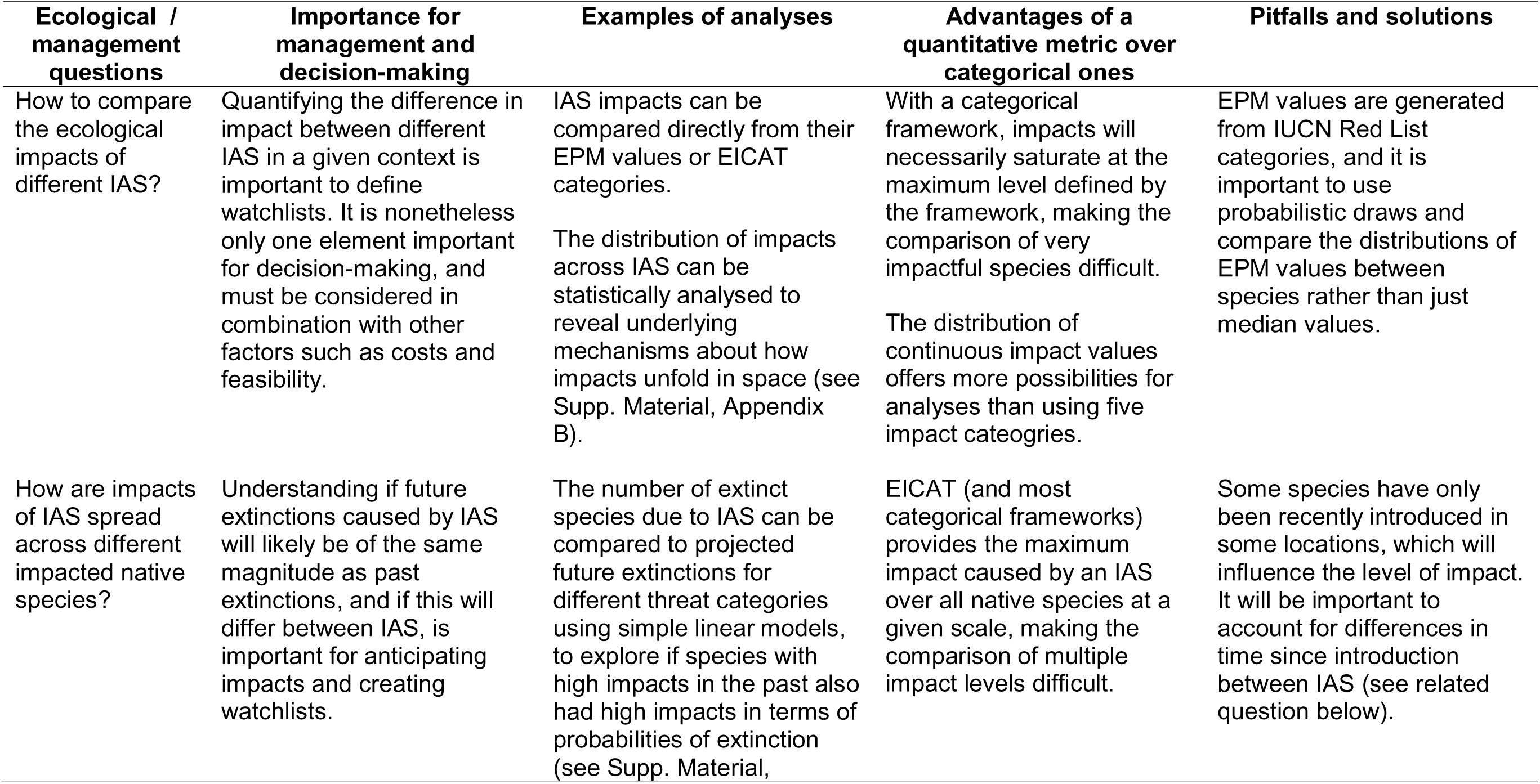
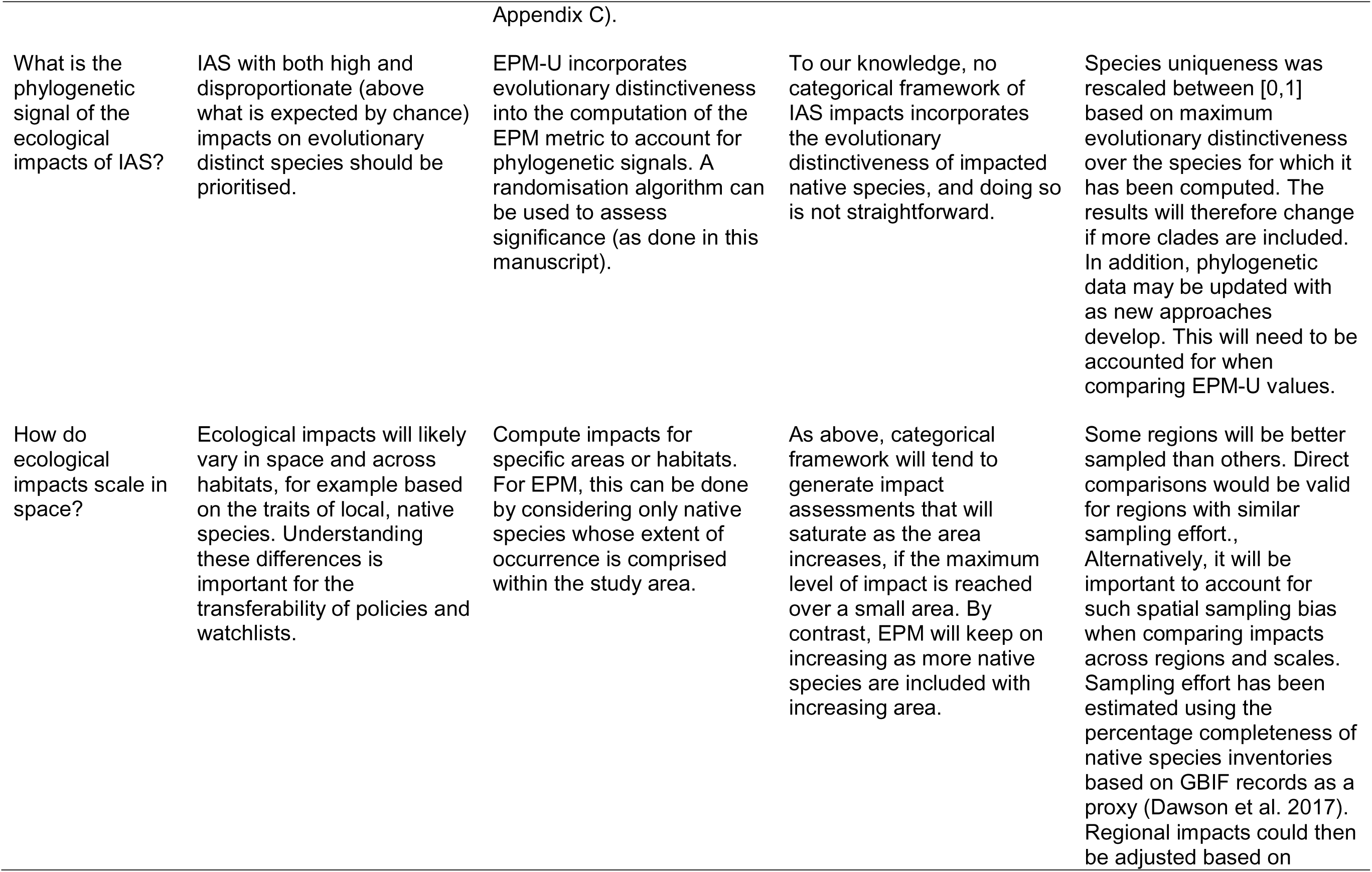
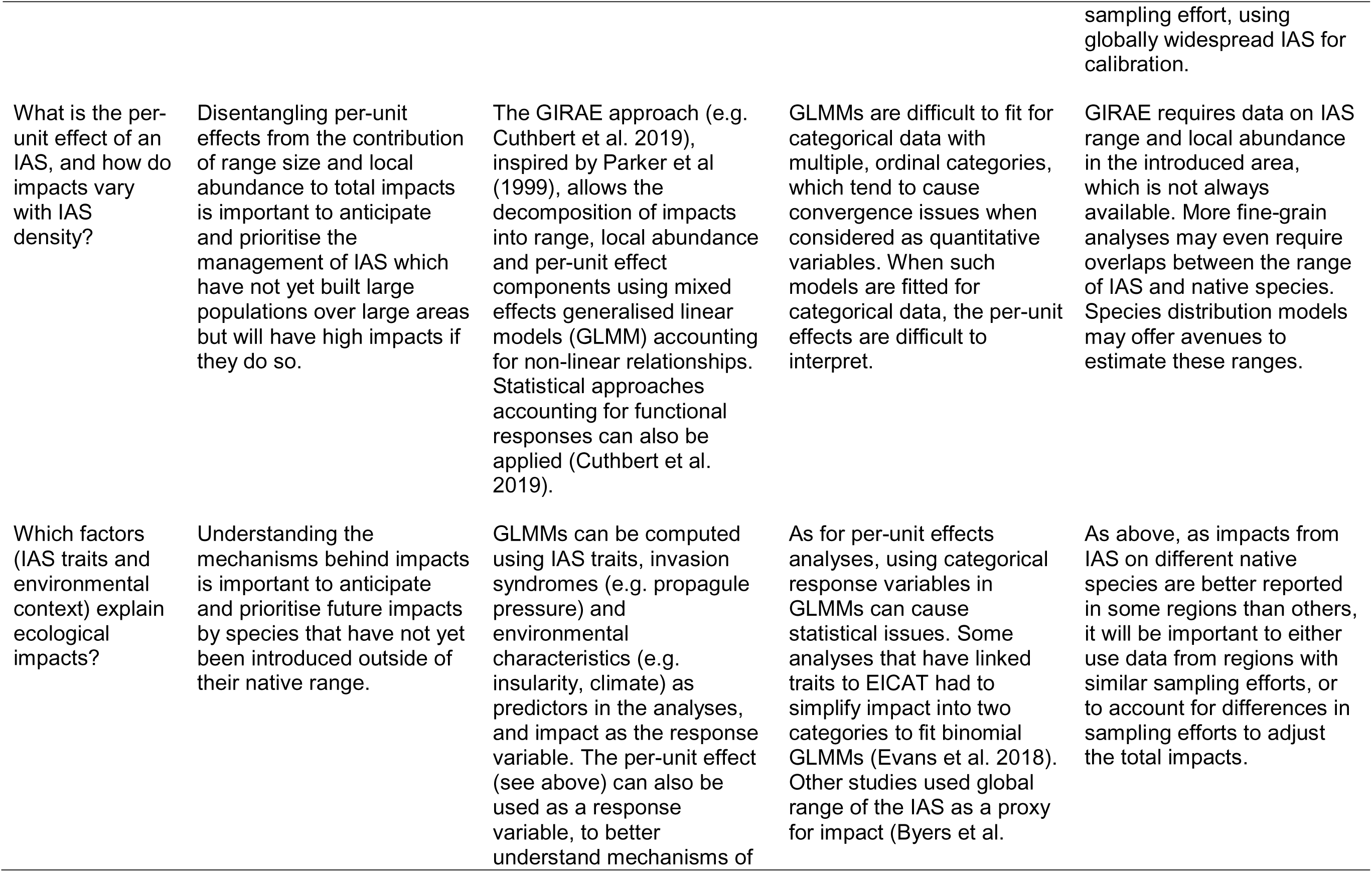
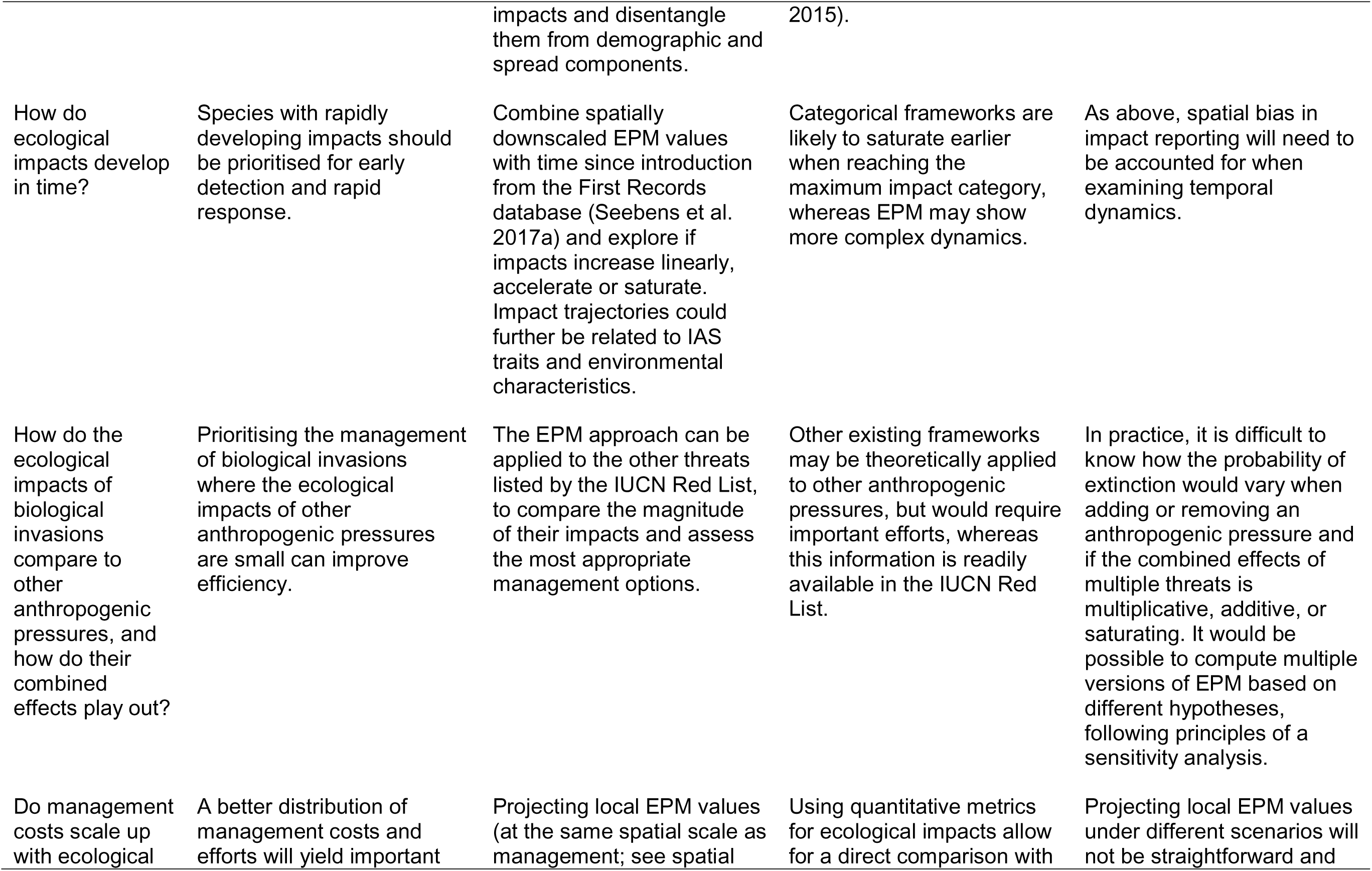
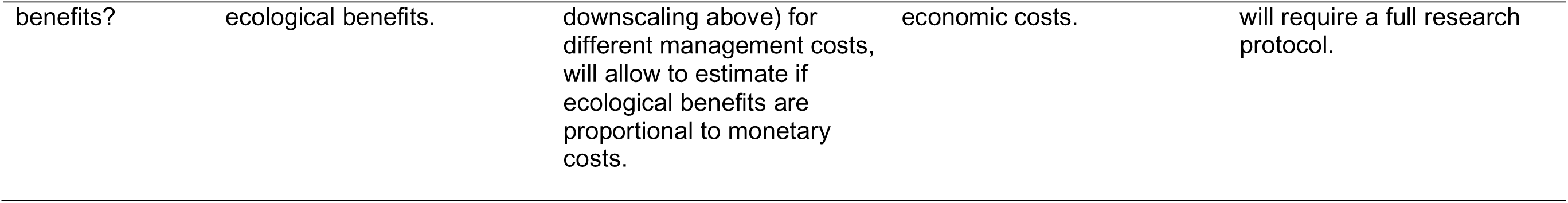
Non-exhaustive list of questions relative to the ecological impacts of IAS that can be addressed with EPM, along with the importance of answering these questions for management and decision-making. We suggest analyses that can be performed to answer these questions and we explain the advantages of using a quantitative rather than a categorical metric to do so. Finally, we indicate some pitfalls for running these analyses, and suggest potential solutions.

Here, we propose the concept of Extinction Potential Metric (EPM) as a measure of impact of individual IAS on native species. EPM uses the existing IUCN Red List of Threatened Species framework (IUCN 2024), converting the Red List category of each native species into a probability of extinction for the whole species within 50 years for each native species threatened by IAS. The EPM score of a given IAS is computed by summing these probabilities over all native species affected by this IAS, resulting in the number of species already and expected to go extinct within 50 years. By doing so, it allows for comparison of IAS effects with different magnitudes on different native species, i.e. affecting their global populations and ranges differently (e.g. species extinction vs. impact on part of the native species global population, which contribute to the risk of extinction of this native species). Translating these different impact magnitudes into probabilities of species extinction enables us to aggregate these impacts into a single metric for multiple native species. EPM thus provides a quantitative synthesis for assessing IAS ecological impacts through harnessing of Red List data, and provides opportunities to answer new ecological questions through different statistical analyses (Table 1).

We introduce three specific EPM metrics capturing different aspects of impact. First, the absolute EPM (EPM-A) of an IAS can be conceptualised as the number of species already extinct that were threatened by this IAS, plus those that are threatened by this IAS and are expected to go extinct within a 50-year time frame under a business-as-usual scenario. Second, the relative EPM (EPM-R) assesses the relative impact of IAS after accounting for other anthropogenic pressures listed by the IUCN Red List (such as agriculture and aquaculture, biological resource use, or climate change and sever weather). Finally, as IAS often threaten specific families and functions, especially on islands (Sayol et al. 2021; Soares et al. 2021; Marino et al. 2022), we introduce EPM-U, which represents the number of ‘unique’ native species that have gone and would go extinct within a 50-year time frame, where species uniqueness accounts for the evolutionary history of the impacted native species, following the EDGE2 framework (Gumbs et al. 2023). Below, we present the EPM-A, EPM-R, and EPM-U metrics (referred to collectively as EPM) and their computations in detail. We showcase these metrics with a proof of concept on native terrestrial vertebrates (amphibians, reptiles, birds, and mammals) threatened by IAS. We focus on native terrestrial vertebrates because the IUCN Red List has evaluated over 80% of described species for vertebrates, whereas only 18%, 2% and 0.8% of plants, invertebrates and fungi/protists have been evaluated, respectively (IUCN 2024). Fishes were excluded because the use of the Red List for marine species has been questioned (Ojaveer et al. 2025). We also show how EPM can be used to identify which impact mechanisms are most detrimental to native species (such as species mortality or competition; IUCN and CMP 2012a). We also showcase the potential of EPM for improving our understanding of IAS impacts by providing preliminary quantitative analyses comparing impact value distributions across taxa and by disentangling past and projected future extinctions (Supp. Material Appendices B & C) and suggest a list of ecological questions it will contribute to answer in the future (Table 1). We nonetheless discuss the limitations of EPM stemming from existing biases in the IUCN Red List, including the role of expert opinion to assign Red List categories to species and associated threats. By providing a quantitative synthesis of known ecological impacts of IAS recorded in the IUCN Red List, EPM advances our understanding and assessment of IAS impacts. EPM complements other categorical metrics that provide additional information, including reversibility of impacts and different spatial scales, or which can be more readily applied to decision-making, in particular EICAT (Van der Colff et al. 2020).

## Method

### Data: Native species impacted by identified invasive alien species

The IUCN Red List applies a set of standardized, quantitative criteria based on species’ global population size, trends, and geographic distribution, which are interpreted by expert assessors to assign extinction risk categories. We use the Red List as a globally recognized and standardized dataset that remains foundational for biodiversity assessments and conservation planning, although we acknowledge the limitations and criticisms of the IUCN Red List and its reliance of expert opinion (see the Discussion for more details on these limitations). More precisely, we utilised the IUCN Red List database (IUCN 2024) to catalogue terrestrial vertebrate species impacted by IAS, i.e. those associated with Threat 8.1 (Invasive non-native/alien species/diseases). We identified 2178 amphibians, 920 birds, 865 reptiles, and 473 mammals in categories Least Concern (LC), Near-Threatened (NT), Vulnerable (VU), Critically Endangered (CR), Extinct in the Wild (EW), and Extinct (EX) for which at least one IAS was listed as a threat (Fig. 1, step 1). The 196 invasive organisms threatening these native species were identified at the species level when possible (73%), but some were only identified at the genus (7%), family (1%), or order and class (0.6%) level. For the remaining threats (19%), the invasive organism was not identified and was recorded as “Unspecified species”. We also distinguished between native species endemic to islands and other species, using data from Marino et al. (2022). Most reported impacted species were native to the Americas (43.2%), followed by Oceania (26.2%), Asia (12.5%), Africa (12.3%), Europe (5.2%) and Antarctica (0.6%) (Supp. Figure A1).

**Figure 1.**
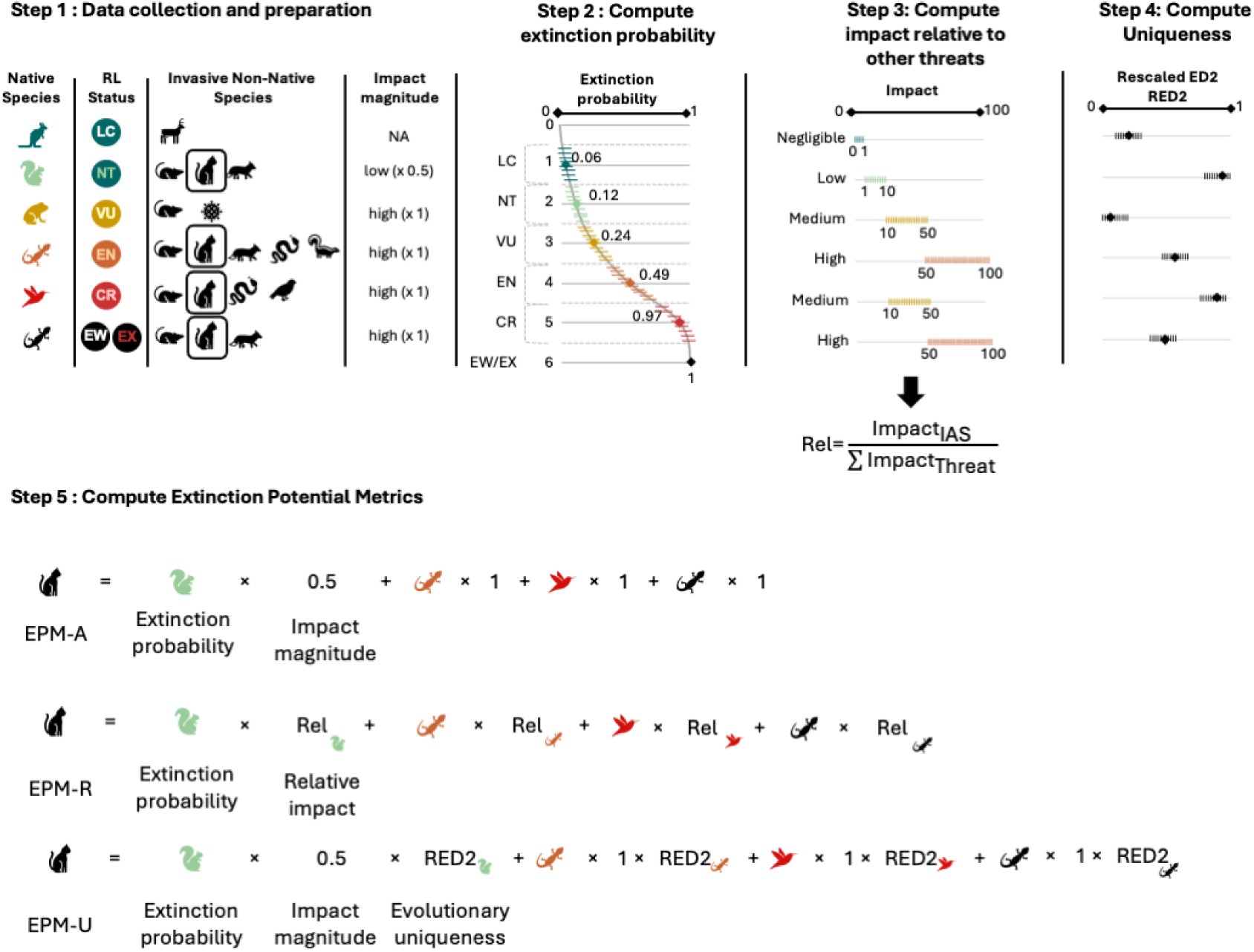
Workflow used to compute the Extinction Potential Metric. Coloured dashes (step 2) depict random draws along the quartic function for extinction probabilities, in the range of indicated values for relative impacts, and between lower and upper interquartiles from Gumbs et al. (2023) for ED2 scores. Relative impact of an IAS on a native species compared to all anthropogenic pressures (step 3) was computed following the approach of Sandvik and Pedersen (2023) and Garnett et al. (2019), which first computes an impact score for each threat affecting the native species using the distributions, and then computes the relative impact of IAS using a ratio. Species uniqueness (step 4) was computed by rescaling ED2 scores from Gumbs et al. (2023); although they are indicated for all species here to show the general principle, ED2 scores were only available for native mammals. For impact magnitude, NAs were either considered as no impact (x 0) or randomly drawn as low or high impacts.

In addition to tabular information summarising the threat details on native species described above, the IUCN Red List provides more detailed textual assessments of the threats affecting native species, including from IAS, interlinking data with the Global Invasive Species Database implemented by the IUCN Invasive Species Specialist Group. Examination of these textual descriptions revealed that species included in the description are often missing from the tabular information. We manually examined the description for the 473 mammals considered, and generated an extended database including all species mentioned both in the text and in the table. Due to the time-consuming nature of manually examining such a large amount of data, this was only done for mammals in the context of the present study, to illustrate the need for data curation in the IUCN Red List.

### Computing the impact magnitude of IAS on native species

To assess the varying impact magnitudes of IAS, we derived data on the scope (proportion of the global population affected) and severity (rate of global population decline) of threats from the database. Consistent with the approaches first proposed by Marino & Bellard (2023) and Marino et al. (2024), species experiencing over 50% global population decline at rates described as very rapid (>30% over 10 years or three generations, whichever is longer), rapid (20–30% over the same period), or slow yet significant (<20% over the same period) due to IAS, were classified as facing high impact and those not meeting these criteria were deemed to have low impact (Fig. 1, step 1). If data on extent or severity were unavailable, the impact was labelled as ‘not available’ (NA) (68 % of the assessments). The distribution of impact magnitudes across taxa is shown in the supplementary material (Supp. Fig A2).

### Computing the probability of extinction of each native species

We calculated probabilities of extinction within 50 years for each native species using the methodology of Marino et al. (2024), inspired by the EDGE2 approach (for Evolutionarily Distinct and Globally Endangered; Gumbs et al. 2023). Probabilities of extinction within 50 years were assigned to each IUCN Red List category as follows: 0.06 for LC, 0.12 for NT, 0.24 for VU, 0.49 for EN, 0.97 for CR, as defined by Gumbs et al. (2023). Extinct in the wild (EW) and Extinct (EX) species were assigned a probability of 1. Although some global policies are set over shorter time frames (for example, the Kunming-Montreal GBF set targets for 2030 and 2050), probabilities of extinction over shorter time horizons are not available to our knowledge. In addition, the shorter the time horizon, the more uncertain these probabilities would become for low-threat categories. To represent the fact that probabilities of extinction should increase continuously with species rarity rather than jump from one category to another, which could generate false precision, a quartic distribution was fitted between IUCN categories (converted to ordinal values from 1 for LC to 5 for CR) and these probabilities of extinction (using the ‘polyfit’ function of the *pracma* R package; Borchers, 2023; Fig. 1, step 2). As EX and EW species were assigned a unique probability of extinction of 1, they were not included in the quartic distribution. To incorporate uncertainty across this quartic relationship, for each species, ten extinction probabilities were randomly generated. A random value was first drawn from a truncated uniform distribution between [-0.5,0.5], and added to the ordinal value corresponding to their IUCN category (i.e. generating a value between [0.5,1.5] for LC, [1.5,2.5] for NT, etc.; Figure 1) using the ‘runif()’ R function. Each of these values was then transformed into a probability of extinction using the quartic distribution (Figure 1; unreported results for 20 and 50 random draws yields similar results, with an ANOVA yielding a *p*-value = 0.98).

### Translating the extinction risks of native species into future number of species extinct due to IAS

Whereas Marino et al. (2024) focused on assessing which native species are impacted by invasive alien species (IAS), our approach shifts the perspective to the IAS themselves. We calculated the absolute extinction potential of each IAS as the sum of the extinction probabilities of all native species it affects, weighted by the severity of its impact (Fig. 1, Step 5, EPM-A). This method highlights not who is affected, but which IAS pose the greatest threat. Extinction probabilities were divided by 2 for low impacts and undivided for high impacts (a factor 2 was chosen as 50% of the global population had to be affected for impact magnitude to be considered as high—a high impact magnitude therefore corresponds to affecting roughly twice the population affected by a low impact magnitude). For each IAS, we calculated 10 EPM-A values, corresponding to the ten extinction probabilities determined per native species. This was done for all native species and for each taxonomic group separately. We accounted for data deficiency in impact magnitude by computing EPM-A values in two different ways. First, following a conservative approach, we excluded threats whose impact magnitude was classified as “not available” (NA), under the presumption that they might represent negligible impacts. Second, acknowledging that NAs may not signify an absence of impact but rather likely reflect gaps in data reporting, we also computed EPM-A scores after randomly assigning a low or high impact to NA threats. This was performed 100 times to address the uncertainties associated with these data gaps, leading to the computation of 1000 weighted probabilities of extinction for the native species, derived from multiplying the 10 extinction probabilities by 100 random weight assignments.

To illustrate the advantages of expressing originally categorical ecological impacts in a quantitative fashion, we performed two sets of statistical analyses, described in more details in the Supplementary Material. First, we modelled the empirical distribution of EPM-A values for each taxonomic group to examine if these differed between taxonomic groups, potentially revealing different underlying processes (Supp. material Appendix B). Second, we explored the relationships between past, imminent and future extinctions by comparing EPM-A values including (i) native species in the EX and EW Red List categories only (i.e. the number of native species that went extinct because of the IAS; EPM-A-Past hereafter), (ii) EPM-A for native species in the CR category (i.e. native species with a high probability of extinction; EPM-A-High hereafter) and (iii) EPM-A for native species in the EN, VU, NT and LC categories (i.e. native species with a lower probability of extinction; EPM-A-Low hereafter). We compared these three EPM-A values for all species, and we assessed if species with high impacts in the past will also have high impacts in the future, and if these relationships are linear (Supp. material, Appendix C).

### Considering unspecified IAS

As some IAS were only identified at the genus and family levels in the IUCN Red List database, we also used information on unspecified species to compute the maximum possible EPM-A value for an IAS based on available data, by assuming unspecified species included all species in the reported genus or family. In other words, we considered a given IAS to be responsible for all the threats for which it is identified, but also for all the threats imposed by unspecified IAS of the same genus or family. For example, the maximum EPM-A score of *Rattus rattus* was computed by aggregating the EPM-A score of *R. rattus*, the EPM-A score of Unspecified Rodentia and the EPM-A score of Unspecified *Rattus*. The maximum EPM-A score of the other *Rattus* species, i.e. *R. norvegicus* and *R. exulans*, were computed in the same way. Conversely, doing so implies that native species impacted by unspecified *Rattus* species were considered as affected by all *Rattus* species (i.e. *R. rattus*, *R. norvegicus*, and *R. exulans*), and will therefore lead to double-counting when all species are not co-occurring over the range of the native species, and an overestimation of impact based on available information for these absent IAS. This maximum score can therefore be seen as an upper limit for EPM-A based on currently available data (but EPM-A scores are still likely underestimated given that not all impacted native species are reported in the IUCN Red List). The four parametric distributions were also fitted to these EPM-A values.

### Accounting for other anthropogenic threats

In practice, many native species are impacted by multiple anthropogenic threats besides biological invasions (1225 out of the 2178 species). Although it is difficult to accurately disentangle the relative impacts of different threats on native species, we propose an approach that assesses the relative impact of IAS while accounting for other threats, assuming these impacts are additive. We followed the approach of Garnett et al. (2019) and Sandvik and Pedersen (2023), which classifies the impact of an anthropogenic threat on a given native species based on its severity and scope as high, medium, low, negligible, or ‘not available’ (NA) (Supp. Table A1). After excluding threats with NA impacts (71% of the assessments), a uniform distribution of impact magnitudes was defined for each category as follows: high = [100, 50], medium = [50, 10], low = [10, 1], and negligible = [1, 0], and a value was randomly drawn in the corresponding distribution for each threat. These impact magnitudes were computed independently for each anthropogenic threat. The relative impact of IAS on a native species was then computed as the ratio of impact value for biological invasions to the sum of impacts values for all threats affecting the native species, therefore generating a value in [0,1] (Fig. 1, step 3). This was done 10 times for different random draws per native species. EPM scores accounting for the relative impacts of different anthropogenic threats (hereafter EPM-R) were then computed using these relative impacts as weights for each extinction probability (Fig. 1, step 5 EPM-R).

### Accounting for evolutionary distinctiveness

Evolutionary distinctiveness is an important component of biodiversity that can be incorporated into EPM by following an approach similar to that used to compute the EDGE2 metric (Gumbs et al. 2023). The EDGE2 score of a species represents the number of millions of years of evolutionary history lost if a species goes extinct. It is the product of the heightened evolutionary distinctiveness (ED2) of a species and of its risk of extinction (GE2, for global endangerment). ED2 accounts for the phylogenetic distance of a species from other species (its “raw” evolutionary distinctiveness ED), but also the extinction risk of these other species. A species with an intermediate ED whose phylogenetically related species are endangered will therefore have a higher ED2, as the risk of losing the whole clade is then higher.

To account for evolutionary history in measuring the impact of IAS, we propose an additional metric, the Extinction Potential Metric for Unique species due to an IAS (EPM-U), which incorporates ED2 scores into EPM. As ED2 values are only available for mammals (Gumbs et al. 2023), we only computed EPM-U of IAS on this taxonomic group. First, we generated a “uniqueness score” ∈ [0,1] for each impacted native species (Fig. 1, step 4). To do so, we rescaled ED2 values by dividing them by the maximum ED2 value, which indicates the most unique species in the database. As Gumbs et al. (2023) account for uncertainty in the calculations of ED2 and report medians and interquartile ranges for each species, for each native species, we randomly sampled 10 ED2 values from a normal distribution where the mean is the median ED2 value and the standard deviation is the lower interquartile interval divided by 1.35. These 10 ED2 values were rescaled between 0 and 1 as described above. Following the same methods as for EPM-A, we then generated 100 scores per IAS.

In addition, we performed a permutation test to examine if IAS-affected species that were more or less evolutionarily distinct than expected by chance. To do so, we randomised the ED2 scores of all native mammals 999 times, and recomputed the EPM-U scores of IAS following the procedure described above, i.e. 100 scores for each permutation. We then computed the median of the 100 EPM-U scores for the real and the 999 randomised ED2 scores, and we computed the 5% and 95% quantiles of these 1000 EPM-U scores. A median EPM-U for the real ED2 score above the 95% quantile would indicate that the IAS affects species that are more evolutionarily distinct than expected by chance, and vice versa (we used a 5% and a 95% threshold because we considered these as two distinct one-tailed tests). The permutation test was performed for the raw data extracted from the IUCN Red List including unspecified species, and for the transformed data for which we considered a native species to be affected by all species in the genus or family when an unspecified species was reported, as described above.

### Linking impact strength to impact mechanisms

The IUCN Red List lists 12 impact mechanisms, with associated definitions (IUCN and CMP 2012a), through which IAS affect native species, using EPM-A values: ecosystem conversion, ecosystem degradation, indirect ecosystem effects, species mortality, species disturbance, hybridisation, competition, loss of mutualism, loss of pollinator, inbreeding, skewed sex ratios, and reduced reproductive success. As for any other part of the Red List assessments, these impacts are identified based on best available data (i.e. to our knowledge, a mix of published data, grey literature and expert knowledge), with a narrative justification and references. To assess the importance of each of these 12 mechanisms, we extracted the native species impacted by any IAS (identified or not). We then computed the corresponding EPM-A score based on the IUCN Red List category of the native species, the severity, and the scope of the threats, generating an EPM-A score for each impact mechanism following the same framework as detailed above. This was done for all native species and for each taxonomic group separately. As for IAS, the distribution of impact magnitudes across mechanisms is shown in the supplementary material (Supp Fig A3).

## Results

### Global impacts of IAS on native taxa

Across all native taxa, median EPM-A values ranged from 0.02 (*Pavo cristatus*, Indian peafowl) to 89.8 (*Batrachochytrium dendrobatidis*, amphibian chytrid fungus) extinct species, with a mean value of 2.4±9.6, when excluding native species with an impact level defined as ‘NA’ (all EPM-A values hereafter correspond to median values). EPM-A values followed a skewed distribution, with a few species having high values and most having low values; 47 of the 196 IAS had a value above one extinct species (Fig. 2.A). The number of species in the different IUCN Red List categories, and therefore with different probabilities of extinctions, varied across the different IAS (Supp. Figs A4, A5). When including native species with an impact level defined as ‘NA’ (i.e. randomly assigned to low or high 100 times), 357 IAS were included, and mean EPM-A values increased by 38%, with 80 IAS above one extinct species. The EPM-A scores of IAS with the greatest impact were particularly affected, with *B. dendrobatidis* rising from 89.8 to 380.7, *Felis catus* (domestic cat) from 87.2 to 138.8, and *Batrachochytrium salamandrivorans* (*Bsal*; the pathogen causing chytridiomycosis in frogs) from 15.1 to 88.5 (Supp. Fig. A6.A). However, IAS ranks were not substantially affected as the correlation between EPM-A scores with and without ‘NA’ impacts were strongly correlated (Spearman’s rank correlation coefficient = 0.86, *p*-value < 0.001). This sensitivity analysis indicates that despite the large quantity of data missing for the scope and severity of impacts, relative EPM-A scores are robust estimates. Future developments will nonetheless need to address this data gap to improve robustness.

**Figure 2.**
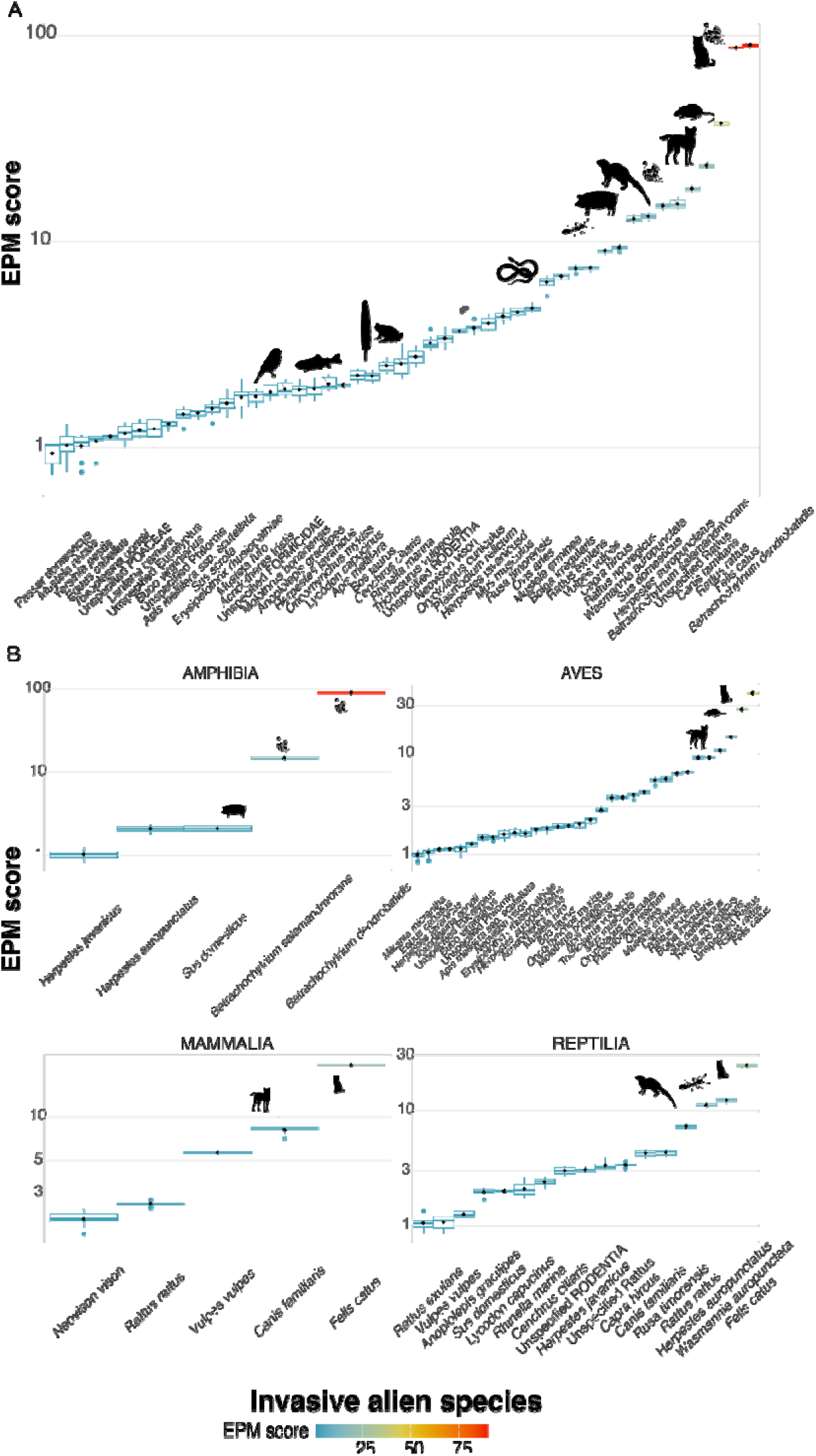
Extinction Potential Metric (EPM-A) scores of invasive alien species (IAS) impacting native terrestrial vertebrates globally using low and high impact assessments only (excluding NAs). Only IAS with a score above 1 are displayed (all taxa: 49 out of 196 species, amphibians: 5 out of 14 species, birds: 32 out of 121 species, mammals: 5 out of 19 species, reptiles: 17 out of 51 species). Panel A shows EPM-A scores based on the impacts of IAS on all native terrestrial vertebrates combined. Panel B shows EPM-A scores based on the impacts of IAS on each taxonomic group (amphibians, birds, mammals, and reptiles) separately. Silhouettes were sourced from PhyloPic.org (T. Michael Keesey, 2025) to illustrate the diversity of invasive alien species (IAS) across the tree of life.

The ranking of IAS differed when considering the different native taxonomic groups separately (Fig. 2.B). Five out of 14 IAS had an EPM-A higher than 1 extinct amphibian species. Two pathogenic fungi were particularly impactful, *B. dendrobatidis* (EPM-A = 89.8), followed by *B. salamandrivorans* (EPM-A = 15.1), with only three other species with an EPM-A value above one. By contrast, 32 out of 121 IAS had an EPM-A above 1 extinct bird species, and three IAS above 10: *F. catus* (EPM-A = 40.1), *Rattus rattus* (black rat; EPM-A = 27.2), and *Canis familiaris* (domestic dog; EPM-A = 10.7) (Fig. 2.B). Five out of 10 IAS had an EPM-A score above one extinct mammal species, *F. catus* (EPM-A = 22.6) being particularly impactful (Fig. 2.B). Finally, 17 out of 51 IAS had an EPM-A score above 1 extinct reptile species, and 3 had a score above 10: *F. catus* (EPM-A = 25.0), *Wasmannia auropunctatus* (electric ant; EPM-A = 12.2), and *Herpestes auropunctatus* (small Indian mongoose; EPM-A = 11.3) (Fig. 2.B). Including native species with an impact defined as ‘NA’ increased EPM-A values, particularly for amphibians (5 to 19 IAS with a score above 1, a 280% increase). The number of IAS above 1 rose by 30% for birds (34 *vs* 26), 100% for mammals (10 *vs* 5) and 60% for reptiles (25 *vs* 15) (Supp. Fig. A6.B). Similarly, we observed substantial increases in EPM-A values for the most impacting IAS for amphibians (*B. dendrobatidis* EPM-A = 380.5 and *B. salamandrivorans* = 87.0) but also for reptiles (*F. catus* EPM-A = 51.8, *R. rattus* EPM-A = 21.3, and *H. auropunctatus* EPM-A = 19.8) and to a lesser extent for birds (*F. catus* with EPM-A = 54.1, *R. rattus* EPM-A = 32.3, and *C. familiaris* EPM-A = 14.4) and mammals (*F. catus* EPM-A = 31.7) (Supp. Fig. A6.B).

Results were similar for maximum EPM-A scores considering unspecified genera or families, with *Rattus* species seeing the main increase, due to many impacts being reported for unspecified *Rattus* and rodents in the IUCN Red List database (Supp. Fig. A7). *Mus musculus* (house mouse) and *H. auropunctatus* had a substantive increase in their score when considering unspecified rodents and unspecified *Herpestes*.

Insular endemic species accounted for 82.0±25.7% of the previously computed EPM-A scores across all native species. Specifically, the impact of IAS was highest for island endemic amphibians and reptiles, with mean changes of −92.4±27.2% and −88.7±16.6%, respectively. The effect was slightly less pronounced in birds, at −80.6±27.8%, and was the weakest in mammals, at −62.0±37.0% (Fig. 3). Adding native species with ‘NA’ impact did not broadly affect these results (Supp. Fig. A9).

**Figure 3.**
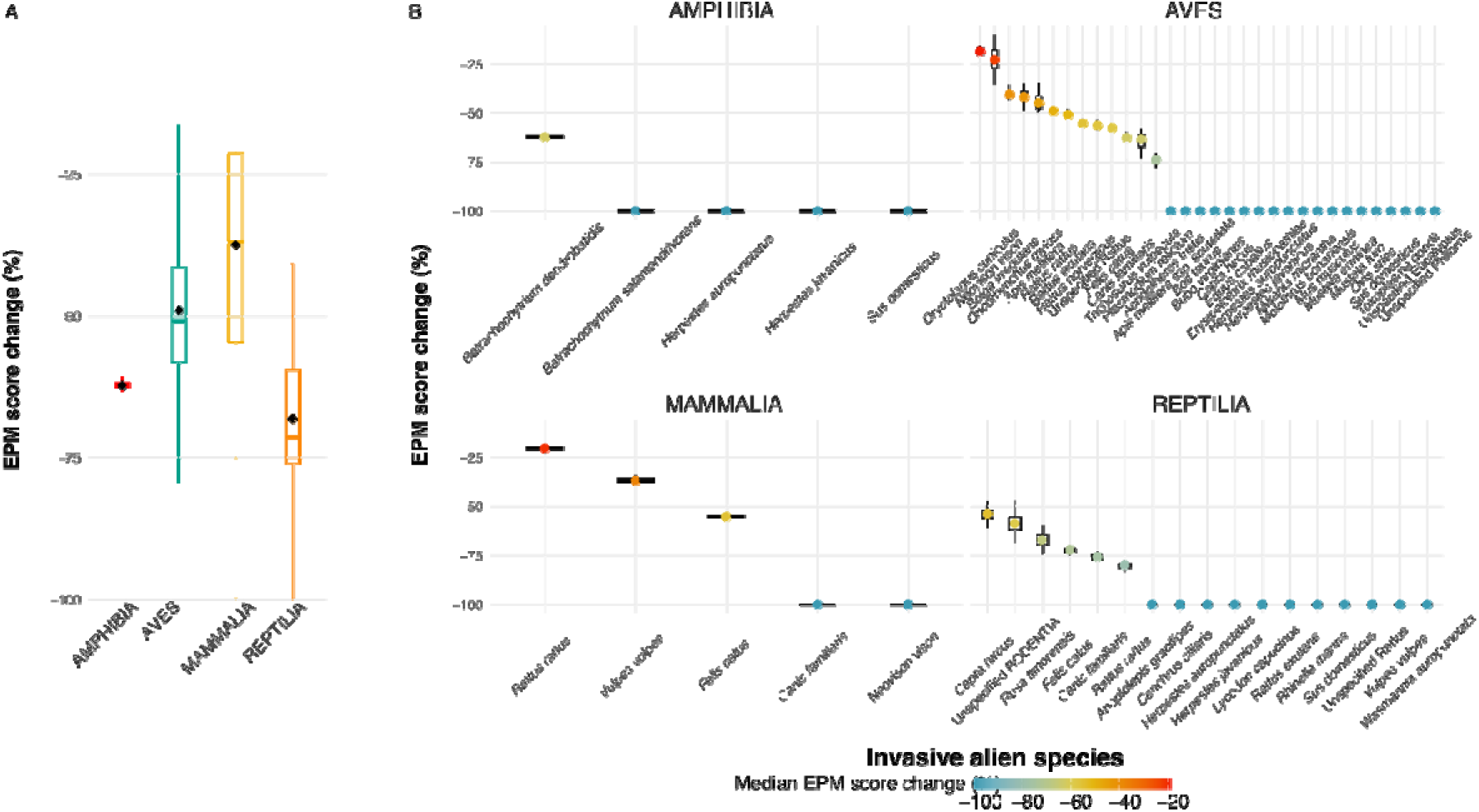
Contribution of insular endemic species to the EPM-A scores. EPM-A scores reflect the impact of IAS on native terrestrial vertebrates globally, considering insular endemic species. Panel A shows the mean change in EPM-A scores (in %) and the associated standard deviation when excluding insular endemic species for each of the four native taxonomic groups. Panel B highlights IAS with a difference in score of more than 50% when excluding insular endemic species.

The distributions of EPM-A values were better fitted by lognormal distributions overall, for all taxonomic groups (Supp. Material Appendix B, Supp. Table B1). When considering only the tail though, power laws provided the best fit. Mammal and reptile distributions had longer tails than amphibians and reptiles.

The comparison of EPM-A-Past, EPM-A-High and EPM-A-Low values revealed that EPM-A-Low > EPM-A-High > EPM-A-Past overall, i.e. future ecological impacts will likely be higher than past ones for most taxonomic groups (Supp. material Appendix C, Supp. Figs C1-4). In addition, the relationships between EPM-A-Past, EPM-A-High and EPM-A-Low were linear, indicating no saturation in impacts (Supp. Table C1, Supp. Fig. C5).

### Impacts of IAS on mammal species using tabular vs textual information and number of unique extinct species

Considering textual information in addition to the tabular data led to include up to 10 more native mammals impacted by a given IAS with a mean value of 0.89±1.12 (Supp. Fig. A10). Including these additional species (with an impact magnitude characterised as ‘NA’) led to 29 IAS having an EPM-A score of more than 1 extinct mammal species (Supp. Fig. A11), i.e. an increase of 24 IAS compared to only using tabular data (Fig. 2). *F. catus* had the largest impacts (EPM-A = 64.2), followed by *C. familiaris* (EPM-A = 30.6), *R. rattus* (EPM-A = 20.0), and *Vulpes vulpes* (red fox) (EPM-A = 19.3).

### Impacts of IAS after controlling for other anthropogenic threats

For native species for which impacts from multiple threats were documented (1225 out of the 2178 native species), EPM-R values ranged from 0.001 (*Lagocephalus sceleratus*, silver-cheeked toadfish) to 84.4 (*B. dendrobatidis*). The average EPM-R was 1.57±7.4, i.e. about half of the average EPM-A score (Supp. Fig. A12.A). EPM-R scores were similar to EPM-A scores for IAS impacting amphibians. By contrast, EPM-R scores were overall about 30% lower than EPM-A scores for birds, mammals, and reptiles (Supp. Fig. A13.A). The ranks of EPM-R and EPM-A scores remained mostly unchanged (*p*-value < 0.001 for Spearman’s rank correlation coefficients for all taxonomic groups, Supp. Fig. A13.A).

### Impacts of IAS on mammal species considering evolutionary distinctiveness

Considering the evolutionary distinctiveness of impacted mammals had little effect on the ranking of the four most impactful IAS, but had important effects on rankings for other IAS, with, for example, *Boiga irregularis* (brown tree snake) losing 95 ranks and *Mucor amphibiorum* (a fungus affecting amphibians) gaining 108 ranks (Supp. Figs A14, A15.B).

Importantly, there were 15 IAS with EPM-U scores lower and 15 IAS with EPM-U scores higher than expected by chance (17 and 13 for EPM-U scores accounting for unspecified species) (Fig. 4, Supp. Fig. A14). These represented more than 5% of the 150 IAS (7.5 IAS). The 15 IAS impacting less evolutionarily distinct native species had EPM-U scores ranked between 21 and 150 out of the 150 IAS (median = 110, median absolute deviation = 49), thus being widespread across all IAS. By contrast, the ranks of the 15 IAS with EPM-U scores above the 95% quantiles ranged between 3 and 61 (median = 20, median absolute deviation = 16), indicating these species were amongst the most impactful.

**Figure 4.**
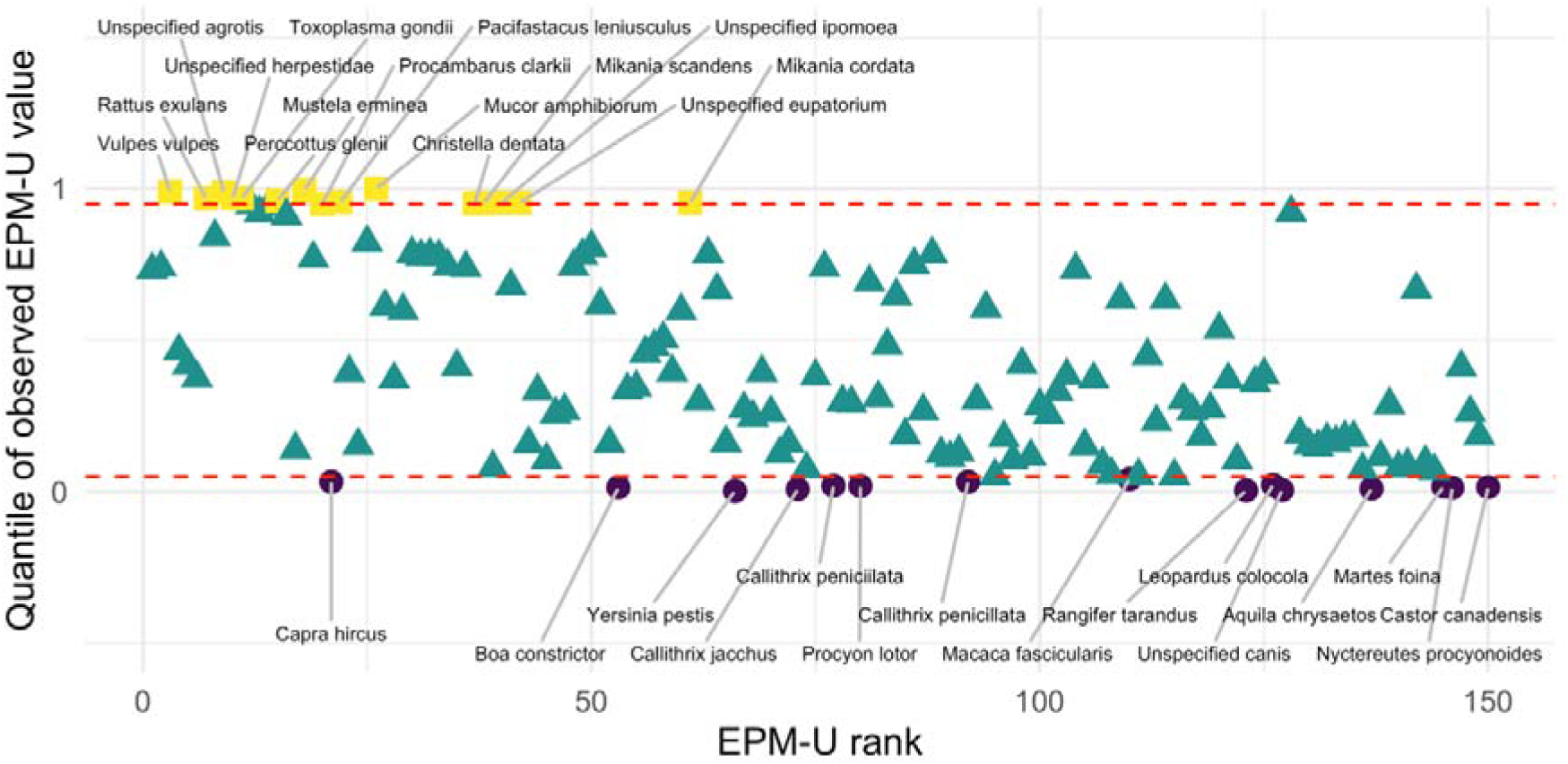
Significance of EPM-U values after randomising ED2 scores 999 times. Low ranks indicate high EPM-U values, and reciprocally. The red dashed lines indicate the 5% and 95% percentile. Yellow squares are species with EPM-U values above the 95% quantile, purple circles are species with EPM-U values below the 5% quantile, and green triangles are species with EPM-U values between these two quantiles.

### Impacts caused through different mechanisms

When considering all taxa, the mechanism ’Species mortality’ was associated with the largest EPM-A value of 396.9, more than three times the EPM-A scores of the second and third threats, namely ’reduced reproductive success’ (EPM-A = 122.5) and ‘ecosystem degradation’ (EPM-A = 109.6) (Fig. 5.A). ’Species mortality’ was also associated with the highest EPM-A scores for amphibians, birds, mammals, and reptiles (Fig. 5.B). For native birds, it was closely followed by ’reduced reproductive success’. For native birds and reptiles, ’ecosystem degradation’ also had a substantial effect (Fig. 5.B). ‘Competition’ ranked second for mammals. Adding native species with ‘NA’ impact did not affect the ranking of ‘species mortality’, which stayed the most impactful mechanism (EPM-A = 1018.7). However, ‘ecosystem degradation’ became the second most important mechanism overall (EPM-A = 235.3), ranking above ’reduced reproductive success’ (EPM-A = 158.4) (Supp. Fig. A16).

**Figure 5.**
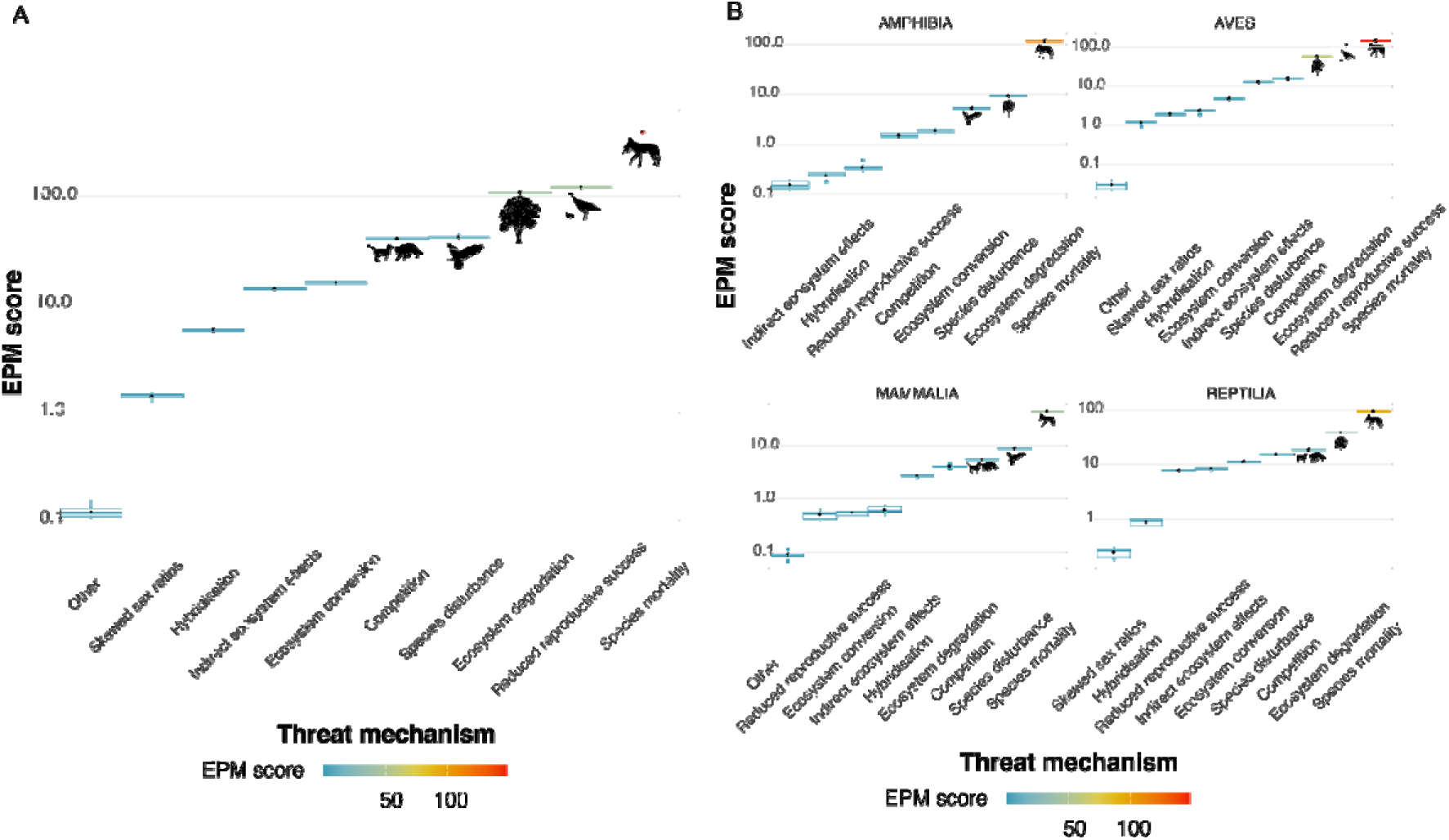
EPM-A scores of impact mechanisms impacting native terrestrial vertebrates globally. Only IAS with a score above 1 are displayed. Panel A shows EPM-A scores based on the impacts of mechanisms on all native terrestrial vertebrates combined. Panel B shows EPM-A scores based on the impacts of mechanisms on each taxonomic group (amphibians, birds, mammals, and reptiles) separately. Silhouettes were sourced from PhyloPic.org (T. Michael Keesey, 2025) to illustrate the most impactful mechanisms.

## Discussion

### EPM is a versatile measure of ecological impacts

The new metric we propose provides a bespoke quantitative synthesis of ecological impact of IAS. The EPM score of an IAS can be interpreted as the number of species that it will have driven to extinction in 50 years under a business-as-usual scenario, including past and future extinctions. EPM therefore uses species as a unit, which is the fundamental unit of conservation, as exemplified by the IUCN Red List of Threatened Species—arguably the most influential global conservation tool to date—on which EPM is based. The IUCN Red List is open access, making the computation of EPM readily applicable and reproducible. As it uses probabilities of extinction, EPM links impacts of different magnitudes, from species extinction to impacts on part of the global native population or range. Being quantitative, it allows more in-depth statistical analyses than categorical metrics, providing finer-scale comparisons and allowing the investigation of underlying processes (Table 1, Supp. Material Appendices B and C). EPM variations allows us to account for other anthropogenic pressures (EPM-R) and for evolutionary uniqueness (EPM-U). Further developments could also include functional distinctiveness or uniqueness, where functional distinctiveness is computed as the average distance between a species and its neighbours in a multidimensional space defined by species functional traits (Marino et al. 2024). Finally, being quantitative, EPM may be compared to other quantitative values important for conservation, such as control costs. If an IAS is more costly to be controlled than another, EPM may reveal if the additional costs would yield proportional ecological benefits (although this will require to compute EPM at the same spatial scale as management costs; see paragraph below of spatial downscaling; Table 1). Thus, EPM will be easily conceptualised by policy-makers and the civil society, and will therefore foster communication about the impacts of biological invasions and the design of management actions, such as native conservation prioritisation.

It is nonetheless important to acknowledge some limitations of EPM. First, although the computation of EPM is transparent and reproducible, the original information provided by the Red List has limitations in terms of transparency and robustness, and is partly based on expert opinion. As a result, the biases present in the IUCN Red List will carry over to EPM, and future work will require careful examination of these biases (see more details on these biases in the context of the present analyses at the end of the Discussion). Second, the quantitative values stem from categorical data (the IUCN Red List categories), which necessarily generates some imprecision in the assessment of the extinction probabilities. To account for this, performing multiple assessments from probabilistic draws allows the generation of EPM value distributions for each IAS. If the mean or median of these distributions is useful for general analyses, it is therefore important to consider the full distributions when comparing specific species, to examine if they are significantly different from each other. This is especially the case when data on scope and severity of impacts are missing: in our analyses, we found a high correlation between median EPM-A scores computed after excluding and including these missing data. Although that indicate that the overall trend in EPM-A scores is robust, the fact that the correlation is below 1 means that the approach will change the ranking for some species when only considering median EPM scores. We therefore recommend examining both sets of scores in addition to the full distribution of each EPM score when comparing specific species, and treat with cautions cases when impact magnitude is shifted between IAS. Finally, rather than using raw EPM values for decision-making (for example via discrete thresholds), we argue that the strength of EPM lies in its versatility (developed in the following paragraphs) and the new analyses this versatility allows, to answer ecological questions important for management, including fine-scale comparisons of impacts across IAS and habitats (Table 1).

EPM offers the possibility to be temporally and spatially downscaled. Regular updates to Red List assessments allow for tracking of trends in extinction risks, as exemplified by the Red List Index (Butchart et al. 2025). It is nonetheless important to acknowledge that updates to Red List assessments are biased towards some taxonomic groups, with birds being historically more frequently updated, whereas the status of many species are rarely re-evaluated (Butchart et al. 2025). This bias would be reflected in temporal trends of EPM for different taxonomic groups. In addition, changes in the status of a species may be due to newly available knowledge or corrections to the use of the available data, rather than an actual change in global population abundance or range (IUCN 2024). This information would therefore need to be incorporated in temporal analyses to document changes to impact at different timescales.

EPM, as presented here, assesses cumulative ecological impacts of IAS at the global scale, because it is the scale at which the IUCN most commonly provides Red List categories and with the most complete level of detail. However, these impacts are likely to vary spatially, especially for globally widespread species. To account for this spatial heterogeneity, as for the Red List Index, EPM can be computed at smaller spatial scales using existing national and regional assessment of Red List categories when available (IUCN 2012; ZSL and IUCN National Red List Working Group 2022). EPM could therefore also be spatially explicit if the same level of information on the status and threats on native species is assessed at multiple spatial scales. This downscaling is particularly pertinent given that the impacts of IAS can differ substantially among populations occurring in different regions over time, undermining the efficacy of global species ‘watch lists’. It would also enable a region to anticipate the impacts of IAS that have not been introduced yet, but have been in other regions with similar climates and biodiversity.

As demonstrated in our analyses, EPM can be computed for different groups of impacted native species, allowing for the capture of multiple dimensions of impacts caused by IAS, rather than generating a single value or category for an IAS, as done by most existing impact classification schemes. Here, we showed that IAS impacts varied across four taxonomic groups of native terrestrial vertebrates. In addition, although we applied EPM at the IAS level, it would be straightforward to upscale the metric to the genus level or beyond, or for any group of IAS. For example, here we also quantified the importance of each mechanism through which IAS impact native species, by computing EPM scores after grouping all IAS impacting native species through these impact mechanisms. Doing so therefore enables us to capture the broader ecological impacts of biological invasions for different components of ecosystems, and how they cascade to *in fine* affect native species survival.

EPM complements existing scoring schemes assessing the impacts of IAS, such as EICAT (Blackburn et al. 2014; Hawkins et al. 2015), not only conceptually, but in also coverage. Interestingly, although 119 IAS are currently characterised by an EICAT impact category in the Global Invasive Species Database (2024) (GISD), there were only 15 species in common with the 196 species extracted from the IUCN Red List. The GISD indicates IUCN Red List species impacted by IAS with an EICAT status. However, many of these IAS are not listed as threats on the Red List, and were therefore not included in our analyses.

Merging information from The Red List and the GISD to compute EPM scores could provide more comprehensive picture of IAS impacts, although merging data from different databases must be done with caution. Moreover, a weak correlation has been observed between the EICAT status of IAS and the Red List status of the native species they impact, and the IUCN Red List and EICAT assessments differ in the geographic scale of the assessment (total native range for the IUCN Red List vs introduced range of an IAS for EICAT), in the type of evidence used in the assessments, in the mechanism of impact classification and in the timing of their updates (Van der Colff et al. 2020). Finally, contrary to EPM, EICAT considers the reversibility of these impacts, and future developments of EPM could use this information. Combining information from EICAT and the IUCN Red List will likely not be straightforward, but would provide a more complete picture of IAS impacts at different spatial and temporal scales (van der Colff et al. 2020).

Native species impacted by biological invasions are often threatened by multiple IAS simultaneously and their effects can combine non-additively. Disentangling the relative impacts of different IAS on a single native species is not possible with the current granularity of IUCN data. It is consequently not possible to assess if an IAS could lead a native species to extinction in the absence of another IAS (e.g. cats are often associated with red foxes in Australia, but have led many native species to extinction by themselves in other regions). In the absence of a clearly identifiable rule to determine the contribution of each IAS, we took a conservative, objective and reproducible approach, attributing the same probability to cause extinction to different IAS affecting a native species with the same level of impact. This follows precautionary principles, an approach already used by other impact classification schemes, including EICAT (Kumschick et al. 2024), to avoid irreversible impacts or high costs to the environment and society that would result from inaction.

Similarly, native species impacted by IAS are also often impacted by other anthropogenic threats (e.g. residential & commercial development, agriculture & aquaculture, pollution, climate change & severe weather, etc., as per the IUCN Red List Threat Classification scheme V3.3; IUCN and CMP 2012b). In practice, it is difficult to know how the probability of extinction would vary when adding or removing an anthropogenic pressure and if the combined effects of multiple threats is multiplicative, additive, or saturating. Here we provide a conservative version of EPM, EPM-R, that accounts for the simultaneous impacts of multiple anthropogenic pressures in an additive fashion. In combination, EPM-A and EPM-R values therefore provide a realistic range of extinction potential values for each IAS.

Anthropogenic threats listed under the IUCN Red List classification scheme have been compared based on the number of native species they impact (Leclerc et al. 2018; e.g. Harfoot et al. 2021). Similarly, EPM could be used to assess the ecological impacts of other anthropogenic pressures individually and in combination. Doing so could lead to a set of harmonised indicators to assess progress towards different targets of the GBF (e.g. Target 7: Reduce Pollution to Levels That Are Not Harmful to Biodiversity; CBD 2022). It could also permit analyses of synergies among global change drivers, by identifying which processes combine to create the highest threats, compared to where they occur individually.

Finally, the versatility of EPM offers avenues to derive indicators assessing the ecological impacts of biological invasions to inform international policy initiatives, in particular Targets 4 and 6 of the GBF. Existing indicators are either lacking or insufficient to track policy performance regarding this target (Henriksen et al. 2024). Such indicators should capture spatial and temporal changes, be species-specific and allow for identification of the mechanisms through which IAS (and other drivers) impact native species (Henriksen et al. 2024), which are all readily included in EPM. Different indicators have been developed, but are either qualitative (e.g. Genovesi et al. 2012), categorical (e.g. Kumschick and Nentwig 2010), do not provide information on the IAS and their impact mechanisms (Butchart et al. 2005; e.g. Butchart 2008), focus on species extinction (i.e. extreme ‘end-points’) rather than combining multiple levels of earlier-stage impacts on different native species (e.g. Bellard et al. 2016a), or consider all threatened and extinct species under a single category (Bellard et al. 2016b). Some studies have incorporated probabilities of extinction and phylogenetic and functional differences, but have focused on impacted species rather than comparing IAS between them (e.g. Bellard et al. 2021). Indicators derived from EPM could be combined with other indicators of impacts (Van der Colff et al. 2020), but also with trends or projections in IAS numbers and ranges, trends in introduction and spread mechanisms, and trends in policy responses (McGeoch et al. 2006, 2010; Rabitsch et al. 2016), to provide a more comprehensive description of the issue and analysis of its drivers. How such indicators will be defined and computed is outside of the scope of this manuscript, as they should be driven by specific conservation questions.

### EPM reveals taxonomic, phylogenetic and temporal patterns of IAS impacts

In our analyses, only a minority of the IAS affecting a taxonomic group had an EPM value above 1. The IAS with the highest EPM-A values (cats [*F. catus*], dogs [*C. familiaris*], black, brown and Polynesian rats [*R. rattus*, *R. norvegicus*, *R. exulans*], foxes [*V. vulpes*], small Indian mongooses [*H. auropunctatus*], and domestic goats [*Capra hircus*], as well as two invasive pathogenic fungi, *B. dendrobatidis* and *B. salamandrivorans* for amphibians; Fig. 2) were, unsurprisingly, already well-known. Most belong to the IUCN list of 100 of the worst invasive species (Lowe et al. 2000; Luque et al. 2014), although there are some noteworthy exceptions, as many species with a high EPM-A value are not on the list (*C. familiaris* being the most important one; Supp. Figure A17). However, as species with small global populations have a higher probability of extinction, an important part of these impacts is recorded for species that are endemic to islands (Fig. 3), confirming the disproportionate impacts IAS have on species that are potentially evolutionarily distinct within insular habitats (Millien 2006), which may be exacerbated by the co-occurrence of multiple IAS on islands (Bellard et al. 2017). As EPM is based on species past and projected future extinctions, and it can reasonably be expected that IAS causing a lot of past extinction have also impacted many species at lower level, it would have indeed been surprising to identify new, previously unrecognised IAS. More importantly, our metrics provide a quantitative framework for comparing the impacts of global invasions in a finer fashion than categorical approaches, including the impacts of species with lower impacts. Thanks to its versatility, EPM will enable new analyses to answer previously unanswered ecological questions (Table 1). For example, fitting the EPM-A scores to four different distributions revealed a better fit with a power law distribution for the upper tail of the distribution, and with a lognormal distribution for the whole data (Supp. Material Appendix B). A powerlZllaw tail means that, once impacts are already large, even larger impacts remain comparatively common. This is consistent with these species impacting native biodiversity in contexts where conditions are different enough for impacts to escalate disproportionately (including prey naivety, or pathogens overcoming epidemiological thresholds). By contrast, the lognormal fit to the whole distribution suggests that moderate impacts result from multiplicative combination of factors (e.g. propagule pressure, habitat suitability, interactions, etc.). However, the distribution fitting, the threshold for the tail of the distribution and the distribution parameters may have also been influenced by biases in how impacts are reported in the IUCN Red List rather than only reflecting ecological processes.

Comparing EPM-A values for EX/EW (past extinctions), CR (species under high level of threat) and EN/VU/NT/LC (species under moderate level of threat) species shows that most impacts in terms of species extinctions over 50 years are yet to come under a business-as-usual scenario (Supp. Material Appendix C). In addition, the linear relationships between EPM-A-Past, EPM-A-High and EPM-A-Low suggests that past impacts provide a good estimate of future impacts. That also suggest that, at the global scale, prevention and management measures of IAS with known impacts have not been sufficient to prevent future ones, while providing further impetus to control existing invasions to mitigate future harm.

These results are nonetheless preliminary and used to illustrate the potential of EPM for further, more comprehensive analyses. Biases in how impacts are reported are likely to influence these results, and future work should, for example, compare reported impacts to historical records on species introductions.

Accounting for other anthropogenic threats when calculating EPM-R shows that the magnitude of the impacts of IAS, compared to other threats, is still substantial, with 31 IAS always obtaining an EPM-R score greater than 1, and with minor impacts on IAS rankings. These high EPM-R scores also imply that management of IAS could yield important benefits for recovering threatened species, particularly at earlier invasion stages before regional impacts escalate for these and closely related species. Such management approaches tend to be relatively inexpensive per unit of area compared to other strategies such as habitat restoration (Reside et al. 2024). Controlling a limited number of IAS could therefore substantially prevent and mitigate impacts of biological invasions on biodiversity, although control may be more efficient when targeting multiple IAS having simultaneous impacts, especially on islands (Glen et al. 2013; Bellard et al. 2017). This corroborates and provides further support to the conclusions of the IPBES assessment that, with sufficient resources, political will and long-term commitment, preventing and controlling IAS will yield significant long-term benefits for nature (IPBES 2023). It is nonetheless important to note that impact is only one factor to consider when deciding which species to manage, as other factors, including costs and feasibility, are also important.

Importantly, considering the evolutionary distinctiveness of impacted native mammals revealed that some of the most impactful IAS disproportionately threatened evolutionarily distinct native species (Fig. 4, Supp. Fig. A15). Evolutionary distinctiveness, and more generally phylogenetic diversity, is a core component of conservation as it captures multiple concepts (Winter et al. 2013; Lean and Maclaurin 2016; Gumbs et al. 2023). Evolutionarily distinct species may be considered to have a high intrinsic or instrumental value on the basis of their uniqueness (Vane-Wright et al. 1991). Phylogenetic diversity (maximised by conserving the most unique species) has also been advanced as a proxy for functional diversity and evolutionary potential, whose maximisation is important for ecosystem services and for maintaining the benefits humans draw from nature, although these claims have been questioned (Winter et al. 2013; Mazel et al. 2018). Nonetheless, phylogenetic diversity remains overlooked in IAS risk assessments. If prioritising IAS for management should also account for management feasibility and costs (Robertson et al. 2021; Kumschick et al. 2025), assessing the risks they pose is an essential component of this process. Here, our results suggest that it may be important to prioritise species with EPM-U scores significantly higher than expected by chance, as these species also tend to be amongst the most impactful IAS. Why some IAS have EPM-U scores higher or lower than expected by chance, and why the most impactful species tend to disproportionately affect evolutionarily unique native species, is unclear and requires further investigation.

“Species mortality” is the main mechanism through which IAS impact native species (Fig. 5), confirming previous assessments based on EICAT (Carneiro et al. 2024). This occurs through predation for mammals, birds, and reptiles, and through two diseases for amphibians. Rankings of the other mechanisms varied substantially across the four taxonomic groups. Interestingly, birds were the only group for which reduced reproductive success was an important mechanism, ranking second. However, this is likely due to the fact that IAS consume native bird eggs before hatching, which is similar to predation as this is linked to direct resource consumption. Aggregating the EPM-A scores of these two impact mechanisms for native birds generates a profile similar to that of the other taxonomic groups. Attention may nonetheless be needed for more subtle impact mechanisms which are less conspicuous than those driving direct mortality, and it is likely that indirect impacts are less frequently reported in the IUCN Red List. In particular, the importance of different mechanisms may change depending on the measure of impact, as it has been shown that ecosystem degradation also had important impacts on phylogenetic and functional diversity in impacted native communities (Bellard et al. 2021).

It is important to understand that the analyses we present here, and the resulting EPM values, represent a first proof of concept for the EPM approach. The IUCN Red List contains substantial gaps and biases regarding how it reports the impacts of biological invasions, which we detail below. As a result, it is likely that future developments of the approach used to compute EPM indices, which will include curation of the data and corrections for biases, will generate different values. The raw EPM values presented here should therefore be taken with caution. Nonetheless, given that we computed distributions of EPM values rather than single values for each IAS, and that IAS rankings are consistent across the different computations of EPM (using two different approaches to account for NAs in impact magnitudes and to consider unspecified IAS), we are confident in the general trends we observe.

Despite being a major conservation resource, numerous criticisms and suggestions for improvement of the IUCN Red List have been raised, particularly regarding data consistency, transparency, and the treatment of multiple threats (e.g., Cazalis et al., 2022). Indeed, we acknowledge that the IUCN Red List is not comprehensive, not fully evidence-based, and not all native or invasive species have been assessed or are up-to-date (Cazalis et al. 2022, 2023). This is exemplified by the large shares of IAS with impacts towards particular native species classified as ‘NA’ in the current study, and the potential spatial bias when reporting IAS as a threat on a native species (Supp. Figure A1). Impact magnitude is poorly informed for terrestrial vertebrates, especially for amphibians, with about 80% of amphibians threatened by IAS having ‘NA’ as impact magnitude (Marino et al. 2024). The resource is also more likely to report impacts from IAS when these have been long standing, and often uses observed and inferred impact through expert opinion rather than comprehensive, published evidence (Van der Colff et al. 2020; Gula et al. 2023). Our analyses were nonetheless designed to account, at least partially, for such imperfect information, by using random draws within probabilistic distributions, resulting in distributions in EPM values. Conversely, despite our results unveiling that IAS with previous impacts can have even more substantial future ones, the IUCN Red List is of limited use in identifying future problematic IAS with a relatively recent invasion history, given time lags to invasion and impacts (Essl et al. 2011). Severity of impacts is estimated over time scales of 10 years or three generations, suggesting that impacts must have occurred over a substantial amount of time to be considered as severe. This could be misleading for IAS that have only recently arrived somewhere, and for which impacts may be important in scope (i.e. affect a large part of the global or regional native species population) and severe (i.e. cause rapid decline in the global or regional abundance of native population) in the future, once its population starts increasing. The figures we present are therefore conservative, and future developments of EPM could consider predicted scope and severity. Doing so will require the development of additional statistical approaches to, first, disentangle the contribution of range size, local abundance and per-unit effects to total impacts (Latombe et al. 2022), and, second, to examine how ecological, functional and phylogenetic data on the IAS and the receiving ecosystems may be used to predict these three components. In addition, high EPM values for a few IAS, particularly those on the 100 of the worst list, could be in part an artefact of increased research and communication efforts which have drawn attention to their impacts. We therefore do not discount that a more comprehensive assessment of other IAS would modify the distribution of impacts, and lead to an overall higher extinction burden.

Finally, although the IUCN Red List data are open access and can easily be extracted from the IUCN Red List website as text files containing data in a tabular format, our further examination of more detailed textual information provided on the webpage of each assessed native mammal revealed that this tabular information was incomplete, with up to 10 IAS missing in the list of threats (Supp. Fig. A10). Computing EPM-A scores for the extended database comprising the textual information revealed that impacts were largely under-estimated, even for the most impactful species, and that ranking differed for many species. As the threats and associated IAS are assessed by different assessors for each native species, there is also a chance that there may be some taxonomic ambiguity, for example for species such as *Sus scrofa* and *Sus domesticus*, as the latter is sometimes considered a subspecies of the former. Furthermore, recent research in marine systems has questioned the use of the Red List, owing to IAS impact assessment gaps and biases, suggesting a need for more targeted studies (Ojaveer et al. 2025). We therefore advocate for an in-depth curation of the IUCN Red List database, to harmonise the information content on IAS threats on native species between tabular and textual information, to ensure consistency across taxa and improve evidence availability. Fostering knowledge on IAS impacts on native species is also paramount to improve these figures and develop a more accurate application of impact metrics such as EPM. We argue that the application of EPM also enables the clear identification of important gaps in the data and will drive further efforts to fill these gaps.

## Conclusion

We introduced the Extinction Potential Metric, a new metric to assess the ecological impacts of IAS. EPM offers a quantitative synthesis of the ecological impacts of IAS using continuous metrics, and provides a fine-grained understanding of those impacts, complementing other existing categorical metrics. EPM computes the equivalent number of native species driven to extinction by an IAS using the IUCN Red List categories to assign probabilities of extinction within 50 years. Doing so enables us to incorporate IAS impacts of different magnitudes over different native species into a single metric, and also to account for evolutionary distinctiveness between impacted native species, and for other anthropogenic changes. Applying this metric to native terrestrial vertebrates shows that a few IAS disproportionately affect native species through direct mortality, and that impactful IAS (e.g. *V. vulpes*, *R. exulans* or *Procambarus clarkii*) also often disproportionately affect evolutionarily unique species, suggesting that controlling a limited number of IAS could effectively prevent and mitigate the impacts of biological invasions on biodiversity. If these findings are consistent with existing knowledge and known patterns on IAS impacts, showing methodological reliability, the true potential of EPM goes far beyond reproducing known results. Designed as an open and adaptable framework, EPM can be continuously updated as better or more refined data become available. The metric is versatile, reproducible and transparent, and can be applied to different groups of native species and IAS, and at different spatio-temporal scales, paving the way to unveiling mechanisms of impacts through a range of statistical analyses that would be difficult to run using categorical indices (Table 1). Although the current version is dependent on existing biases from the IUCN Red List, we argue that, in combination with other databases such as the GISD, it can serve as a basis to derive spatio-temporal indicators that will improve our capacity to assess the efficacy of global and regional conservation policies, including for other anthropogenic threats.

## Supporting information

Supplement figures and analyses

## Acknowledgements and funding

We thank Clara Marino and Céline Bellard for useful comments on this manuscript. HS acknowledges funding by the Deutsche Forschungsgemeinschaft (DFG, German Research Foundation) (grant no. 521529463). SK acknowledges the support of the Centre for Invasion Biology (CIB) at Stellenbosch University, the South African Department of Forestry, Fisheries and the Environment (DFFE), and the B3 (Biodiversity Building Blocks for Policy) project, which receives funding from the European Union’s Horizon Europe Research and Innovation Programme (through grant no. 584101059592).

## Data availability

The data on which this paper is based are freely available on the IUCN Red List website (www.redlist.org). A list of the native and invasive alien species used in the analyses is given in the electronic supplementary material.

## Authors contribution

All authors contributed to the study conception and design. Material preparation, data collection and analysis were performed by MPL and GL. The first draft of the manuscript was written by GL and MPL and all authors commented on previous versions of the manuscript. All authors read and approved the final manuscript.

## References

Ahmed DA, Haubrock PJ, Cuthbert RN, et al (2023) Recent advances in availability and synthesis of the economic costs of biological invasions. BioScience 73:560–574. 10.1093/biosci/biad060

Bellard C, Bernery C, Leclerc C (2021) Looming extinctions due to invasive species: Irreversible loss of ecological strategy and evolutionary history. Global Change Biology 27:4967–4979. 10.1111/gcb.15771

Bellard C, Cassey P, Blackburn TM (2016a) Alien species as a driver of recent extinctions. Biology letters 12:20150623–20150623. 10.1098/rsbl.2015.0623

Bellard C, Genovesi P, Jeschke JM (2016b) Global patterns in threats to vertebrates by biological invasions. Proceedings of the Royal Society B: Biological Sciences 283:20152454. 10.1098/rspb.2015.2454

Bellard C, Rysman J-F, Leroy B, et al (2017) A global picture of biological invasion threat on islands. Nature Ecology & Evolution 1:1862–1869. 10.1038/s41559-017-0365-6

Bernardo-Madrid R, González-Moreno P, Gallardo B, et al (2022) Consistency in impact assessments of invasive species is generally high and depends on protocols and impact types. NB 76:163–190. 10.3897/neobiota.76.83028

Blackburn TM, Essl F, Evans T, et al (2014) A unified classification of alien species based on the magnitude of their environmental impacts. PLoS Biology 12:e1001850–e1001850. 10.1371/journal.pbio.1001850

Borchers H (2023) _pracma: Practical Numerical Math Functions_. R package version 2.4.4, <https://CRAN.R-project.org/package=pracma>.

Butchart SHM (2008) Red List Indices to measure the sustainability of species use and impacts of invasive alien species. Bird Conservation International 18:S245–S262. 10.1017/S095927090800035X

Butchart SHM, Akçakaya HR, Berryman AJ, et al (2025) Measuring trends in extinction risk: a review of two decades of development and application of the Red List Index. Philosophical Transactions of the Royal Society B: Biological Sciences 380:20230206. 10.1098/rstb.2023.0206

Butchart SHM, Stattersfield AJ, Baillie J, et al (2005) Using Red List Indices to measure progress towards the 2010 target and beyond. Philosophical Transactions of the Royal Society B: Biological Sciences 360:255–268. 10.1098/rstb.2004.1583

Byers JE, Smith RS, Pringle JM, et al (2015) Invasion Expansion: Time since introduction best predicts global ranges of marine invaders. Scientific Reports 5:12436. 10.1038/srep12436

Carneiro L, Miiller N, Prestes JG, et al (2024) Impacts and mechanisms of biological invasions in global protected areas. Biological Invasions 27:20. 10.1007/s10530-024-03498-w

Cazalis V, Di Marco M, Butchart SH, et al (2022) Bridging the research-implementation gap in IUCN Red List assessments. Trends in Ecology & Evolution 37:359–370

Cazalis V, Santini L, Lucas PM, et al (2023) Prioritizing the reassessment of data-deficient species on the IUCN Red List. Conservation Biology 37:e14139. 10.1111/cobi.14139

CBD (2022) Decision adopted by the conference of the parties to the convention on biological diversity 15/4. Kunming-montreal global biodiversity framework.

Courchamp F, Fournier A, Bellard C, et al (2017) Invasion biology: specific problems and possible solutions. Trends in Ecology & Evolution 32:13–22. 10.1016/j.tree.2016.11.001

Cuthbert RN, Dickey JWE, Coughlan NE, et al (2019) The Functional Response Ratio (FRR): advancing comparative metrics for predicting the ecological impacts of invasive alien species. Biological Invasions 21:2543–2547. 10.1007/s10530-019-02002-z

Dawson W, Moser D, van Kleunen M, et al (2017) Global hotspots and correlates of alien species richness across taxonomic groups. Nature Ecology & Evolution 1:186–186. 10.1038/s41559-017-0186

Diagne C, Leroy B, Gozlan RE, et al (2020) InvaCost, a public database of the economic costs of biological invasions worldwide. Scientific Data 7:1–12. 10.1038/s41597-020-00586-zgas

Diagne C, Leroy B, Vaissière A-C, et al (2021) High and rising economic costs of biological invasions worldwide. Nature 592:571–576. 10.1038/s41586-021-03405-6

Essl F, Dullinger S, Rabitsch W, et al (2011) Socioeconomic legacy yields an invasion debt. Proceedings of the National Academy of Sciences 108:203–207. 10.1073/pnas.1011728108

Evans T, Kumschick S, Şekercioğlu ÇH, Blackburn TM (2018) Identifying the factors that determine the severity and type of alien bird impacts. Diversity and Distributions 24:800–810. 10.1111/ddi.12721

Garnett ST, Butchart SHM, Baker GB, et al (2019) Metrics of progress in the understanding and management of threats to Australian birds. Conservation Biology 33:456–468. 10.1111/cobi.13220

Genovesi P, Carnevali L, Alonzi A, Scalera R (2012) Alien mammals in Europe: updated numbers and trends, and assessment of the effects on biodiversity. Integrative Zoology 7:247–253. 10.1111/j.1749-4877.2012.00309.x

Glen AS, Atkinson R, Campbell KJ, et al (2013) Eradicating multiple invasive species on inhabited islands: the next big step in island restoration? Biological Invasions 15:2589–2603. 10.1007/s10530-013-0495-y

Gula J, Sundar KSG, Willows-Munro S, Downs CT (2023) The state of stork research globally: A systematic review. Biological Conservation 280:109969. 10.1016/j.biocon.2023.109969

Gumbs R, Gray CL, Böhm M, et al (2023) The EDGE2 protocol: Advancing the prioritisation of Evolutionarily Distinct and Globally Endangered species for practical conservation action. PLOS Biology 21:e3001991. 10.1371/journal.pbio.3001991

Harfoot MBJ, Johnston A, Balmford A, et al (2021) Using the IUCN Red List to map threats to terrestrial vertebrates at global scale. Nature Ecology & Evolution 5:1510–1519. 10.1038/s41559-021-01542-9

Hawkins CL, Bacher S, Essl F, et al (2015) Framework and guidelines for implementing the proposed IUCN Environmental Impact Classification for Alien Taxa (EICAT). Diversity and Distributions 21:1360–1363

Henriksen MV, Arlé E, Pili A, et al (2024) Global indicators of the environmental impacts of invasive alien species and their information adequacy. Philosophical Transactions of the Royal Society B: Biological Sciences 379:20230323. 10.1098/rstb.2023.0323

IPBES (2023) Thematic Assessment Report on Invasive Alien Species and their Control of the Intergovernmental Science-Policy Platform on Biodiversity and Ecosystem Services, Roy, H. E., Pauchard, A., Stoett, P., and Renard Truong, T. IPBES secretariat, Bonn, Germany

IUCN (2020a) IUCN EICAT Categories and Criteria. The Environmental Impact Classification for Alien Taxa (EICAT) First edition. IUCN, Gland, Switzerland and Cambridge, UK

IUCN (2020b) Guidelines for using the IUCN Environmental Impact Classification for Alien Taxa (EICAT) Categories and Criteria): First edition. Version 1.1. IUCN, Gland, Switzerland and Cambridge, UK

IUCN (2024) The IUCN Red List of Threatened Species. Version 2024–1.

IUCN (2012) Guidelines for Application of IUCN Red List Criteria at Regional and National Levels: Version 4.0. Gland, Switzerland and Cambridge, UK: IUCN.

IUCN, CMP (2012a) IUCN-CMP Unified Classification of Stresses Version 1.1. Gland, Switzerland and Cambridge, UK

IUCN, CMP (2012b) IUCN-CMP Unified Classification of Direct Threats Version 3.3. Gland, Switzerland and Cambridge, UK

Kumschick S, Bertolino S, Blackburn TM, et al (2024) Using the IUCN Environmental Impact Classification for Alien Taxa to inform decision-making. Conservation Biology 38:e14214. 10.1111/cobi.14214

Kumschick S, Foxcroft LC, Wilson JRU (2025) Advancing the Risk Analysis for Alien Taxa (RAAT) framework. Neobiota 97:319–324. 10.3897/neobiota.97.135975

Kumschick S, Nentwig W (2010) Some alien birds have as severe an impact as the most effectual alien mammals in Europe. Biological Conservation 143:2757–2762. 10.1016/j.biocon.2010.07.023

Langhammer PF, Bull JW, Bicknell JE, et al (2024) The positive impact of conservation action. Science 384:453–458. 10.1126/science.adj6598

Latombe G, Catford JA, Essl F, et al (2022) GIRAE: a generalised approach for linking the total impact of invasion to species’ range, abundance and per-unit effects. Biological Invasions 24:3147–3167. 10.1007/s10530-022-02836-0

Lean C, Maclaurin J (2016) The value of phylogenetic diversity. Biodiversity Conservation and Phylogenetic Systematics: Preserving our evolutionary heritage in an extinction crisis 19–37

Leclerc C, Courchamp F, Bellard C (2018) Insular threat associations within taxa worldwide. Scientific Reports 8:6393. 10.1038/s41598-018-24733-0

Lowe S, Browne M, Boudjelas S, De Poorter M (2000) 100 of the world’s worst invasive alien species: a selection from the global invasive species database. Invasive Species Specialist Group Auckland

Luque GM, Bellard C, Bertelsmeier C, et al (2014) The 100th of the world’s worst invasive alien species. Biological Invasions 16:981–985. 10.1007/s10530-013-0561-5

Marino C, Bellard C (2023) When origin, reproduction ability and diet define the role of birds in invasions. Proceedings of the Royal Society B: Biological Sciences 290:20230196. 10.1098/rspb.2023.0196

Marino C, Leclerc C, Bellard C (2022) Profiling insular vertebrates prone to biological invasions: What makes them vulnerable? Global Change Biology 28:1077–1090. 10.1111/gcb.15941

Marino C, Soares FC, Bellard C (2024) Conservation priorities for functionally unique and specialized terrestrial vertebrates threatened by biological invasions. Conservation Biology 39:e14401. 10.1111/cobi.14401

Mazel F, Pennell MW, Cadotte MW, et al (2018) Prioritizing phylogenetic diversity captures functional diversity unreliably. Nature Communications 9:2888. 10.1038/s41467-018-05126-3

McGeoch MA, Butchart SHM, Spear D, et al (2010) Global indicators of biological invasion: species numbers, biodiversity impact and policy responses. Diversity and Distributions 16:95–108

McGeoch MA, Chown SL, Kalwij JM (2006) A Global Indicator for Biological Invasion. Conservation Biology 20:1635–1646. 10.1111/j.1523-1739.2006.00579.x

Millien V (2006) Morphological Evolution Is Accelerated among Island Mammals. PLOS Biology 4:e321. 10.1371/journal.pbio.0040321

Ojaveer H, Galil B, Seebens H (2025) The commitment of the EU Biodiversity Strategy for 2030 is unfeasible for marine threatened species affected by invasive alien species. Marine Policy 173:106582. 10.1016/j.marpol.2024.106582

Pagad S, Genovesi P, Carnevali L, et al (2015) IUCN SSC Invasive Species Specialist Group: invasive alien species information management supporting practitioners, policy makers and decision takers. Management of Biological Invasions 6:137–135

Parker IM, Simberloff D, Lonsdale WM, et al (1999) Impact: toward a framework for understanding the ecological effects of invaders. Biological Invasions 1:3–19. 10.1023/A:1010034312781

Rabitsch W, Genovesi P, Scalera R, et al (2016) Developing and testing alien species indicators for Europe. Journal for Nature Conservation 29:89–96. 10.1016/j.jnc.2015.12.001

Reside AE, Carwardine J, Ward M, et al (2024) The cost of recovering Australia’s threatened species. Nature Ecology & Evolution. 10.1038/s41559-024-02617-z

Robertson PA, Mill AC, Adriaens T, et al (2021) Risk Management Assessment Improves the Cost-Effectiveness of Invasive Species Prioritisation. Biology 10:. 10.3390/biology10121320

Sandvik H, Pedersen B (2023) Metrics for quantifying how much different threats contribute to red lists of species and ecosystems. Conservation Biology 37:e14105. 10.1111/cobi.14105

Sayol F, Cooke RSC, Pigot AL, et al (2021) Loss of functional diversity through anthropogenic extinctions of island birds is not offset by biotic invasions. Science Advances 7:eabj5790. 10.1126/sciadv.abj5790

Seebens H, Bacher S, Blackburn TM, et al (2021) Projecting the continental accumulation of alien species through to 2050. Global Change Biology 27:970–982. 10.1111/gcb.15333

Seebens H, Blackburn TM, Dyer EE, et al (2017a) No saturation in the accumulation of alien species worldwide. Nature Communications 8:14435–14435. 10.1038/ncomms14435

Seebens H, Essl F, Blasius B (2017b) The intermediate distance hypothesis of biological invasions. Ecology Letters 20:158–165

Soares FC, Leal AI, Palmeirim JM, de Lima RF (2021) Niche differences may reduce susceptibility to competition between native and non-native birds in oceanic islands. Diversity and Distributions 27:1507–1518. 10.1111/ddi.13298

Van der Colff D, Kumschick S, Foden W, Wilson JRU (2020) Comparing the IUCN’s EICAT and Red List to improve assessments of the impact of biological invasions. NeoBiota 62:509–523. 10.3897/neobiota.62.52623

Vane-Wright RI, Humphries CJ, Williams PH (1991) What to protect?—Systematics and the agony of choice. Biological conservation 55:235–254

Vicente JR, Vaz AS, Roige M, et al (2022) Existing indicators do not adequately monitor progress toward meeting invasive alien species targets. Conservation Letters 15:e12918. 10.1111/conl.12918

Winter M, Devictor V, Schweiger O (2013) Phylogenetic diversity and nature conservation: where are we? Trends in ecology & evolution 28:199–204

ZSL and IUCN National Red List Working Group (2022) National Red List Database. https://www.nationalredlist.org

