## Supplement figures and analyses for "Quantifying extinction potential from invasive alien species"

**APPENDIX A: Supplementary figures and table**

**Supplementary Table A1.** Impact of an anthropogenic threat on a given native species based on its severity and scope as defined by Garnett et al. (2019) and Sandvik and Pedersen (2023).

| **Impact category** | **Criteria** |
| --- | --- |
| High | very rapid decline and whole or majority of extent |
| Medium | very rapid decline and minority of extent  OR  rapid decline and whole or majority of extent  OR  slow, significant decline and whole extent |
| Low | rapid decline and minority of extent  OR  slow, significant decline and majority or minority of extent |
| Negligible | negligible decline and whole to negligible extent  OR  no decline and whole to negligible extent  OR  very rapid to slow, significant decline and negligible severity |
| NA | neither severity nor extent were available |

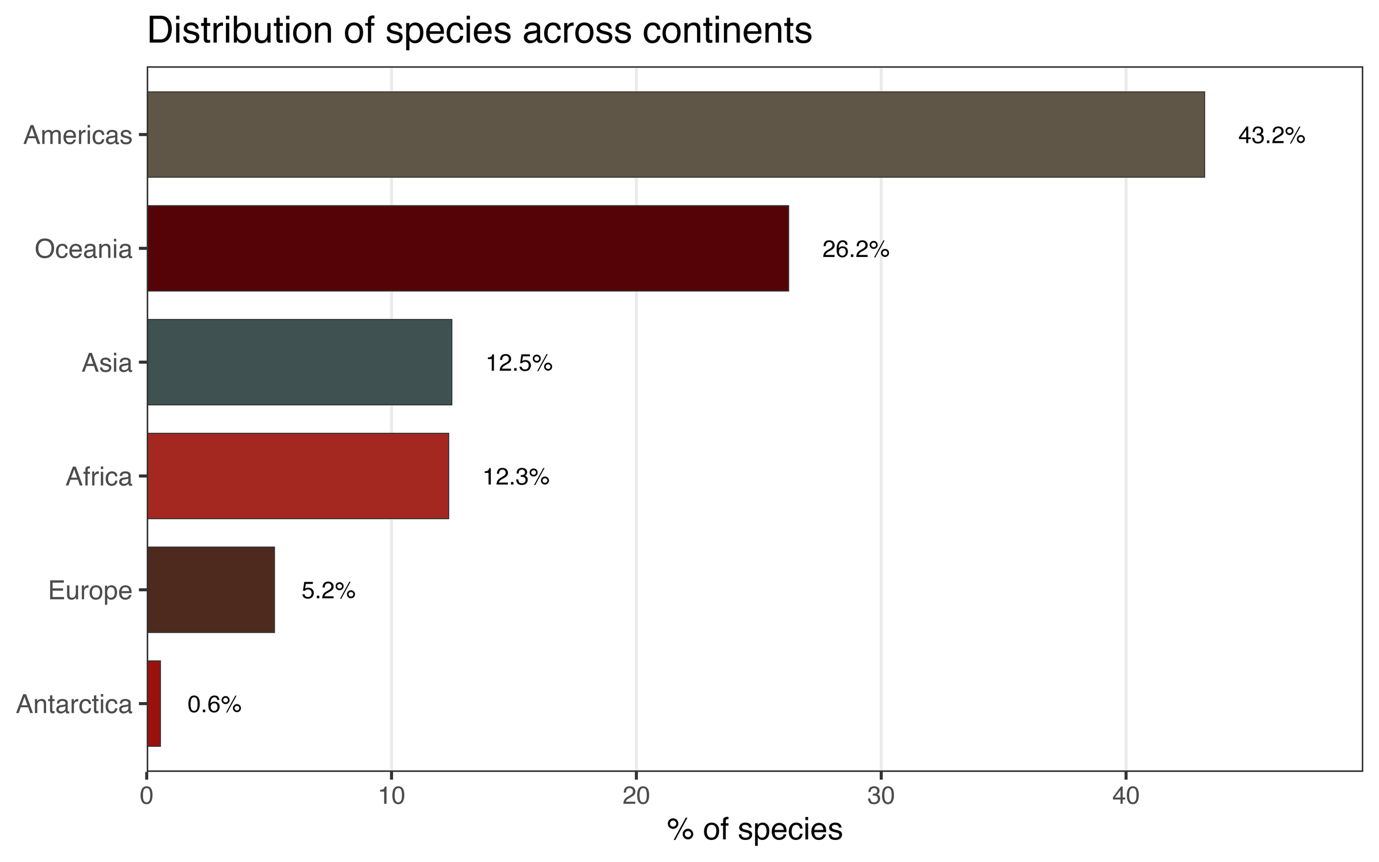

**Figure A1.** Distribution of native species reported to be impacted by IAS on the IUCN Red List at the continent scale.

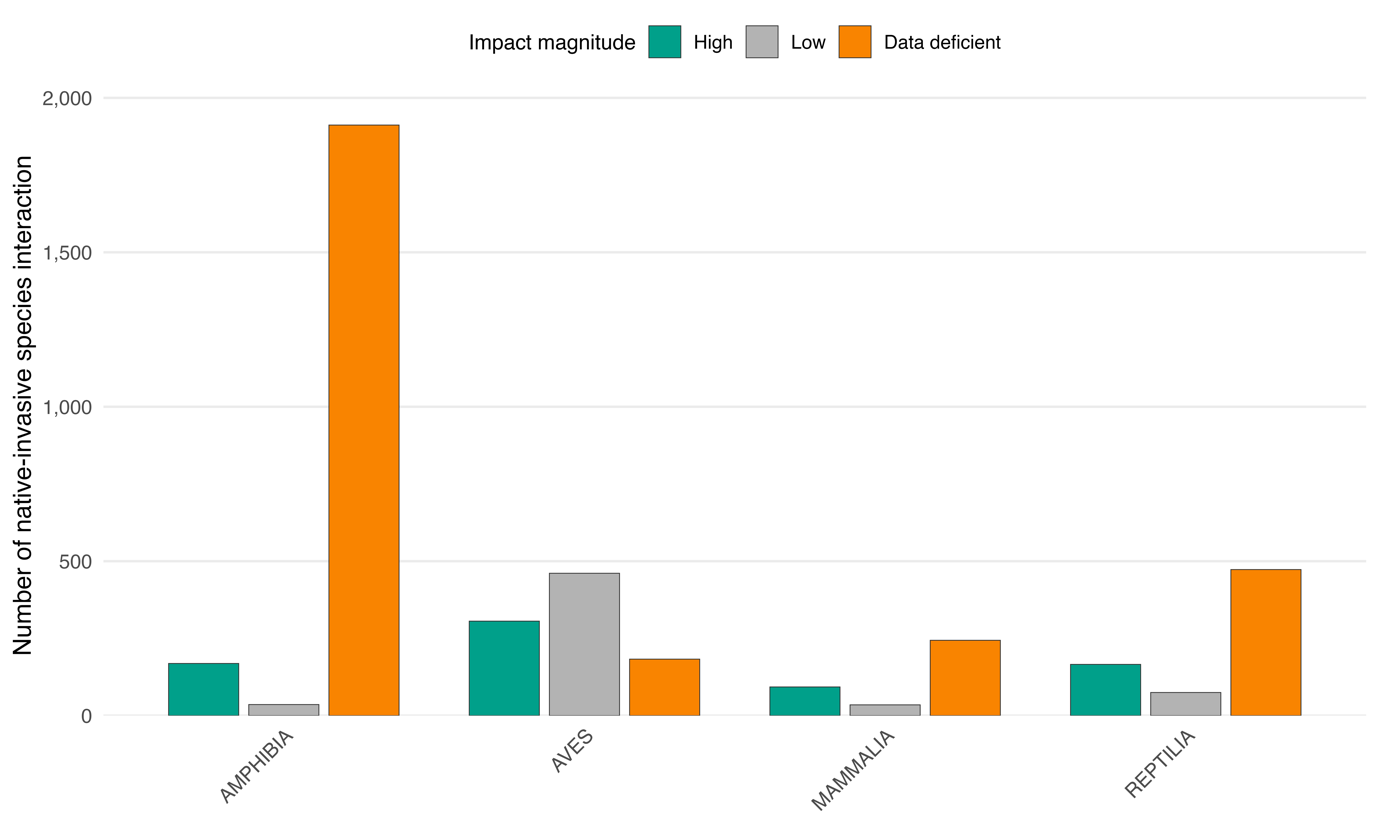

**Supplementary Figure A2.** Distribution of impact magnitudes across taxa. Each impact represents a unique interaction between a native and an invasive species

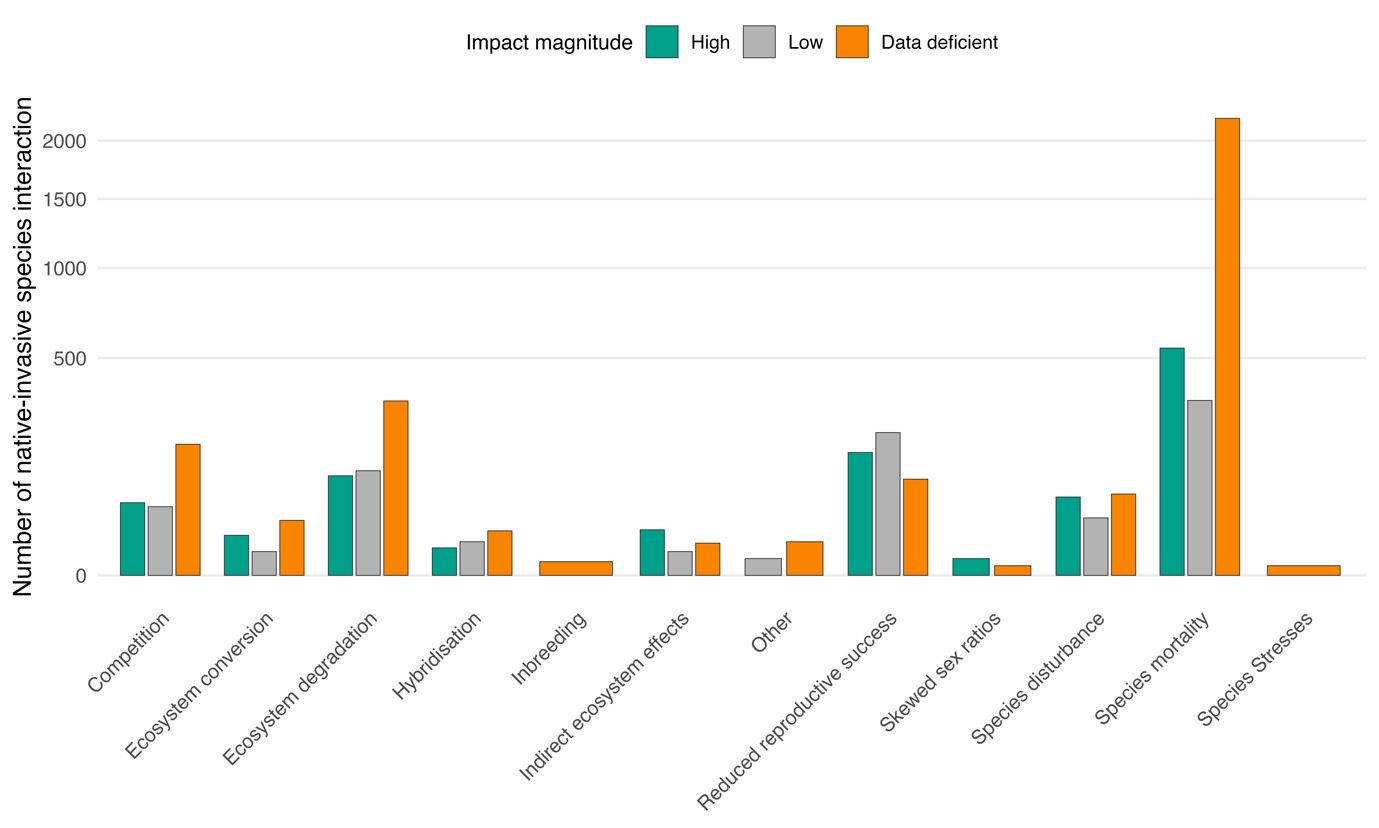

**Supplementary Figure A3.** Distribution of impact magnitudes across impact mechanisms**.** Each data point represents a unique interaction between a native and an invasive species. The y-axis is square root-transformed to improve visualization.

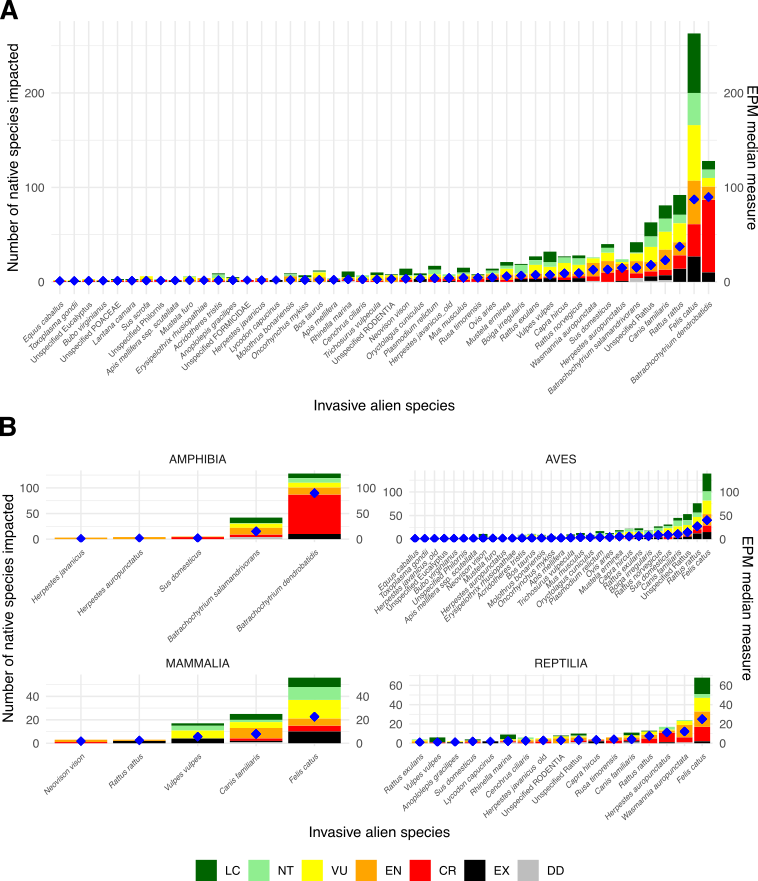

**Supplementary Figure A4. IUCN categories of native terrestrial vertebrates impacted by IAS globally and their associated EPM-A scores using only low and high impact assessments (excluding ‘NA’).** Only IAS with an EPM-A score above 1 are displayed. Panel A shows EPM-A median measure (blue diamonds) and the number of native species per IUCN category, based on the impacts of IAS on all native terrestrial vertebrates combined. Panel B shows EPM-A median measure (blue diamonds) and the number of native species per IUCN category, based on the impacts of IAS on each taxonomic group (amphibians, birds, mammals, and reptiles) separately

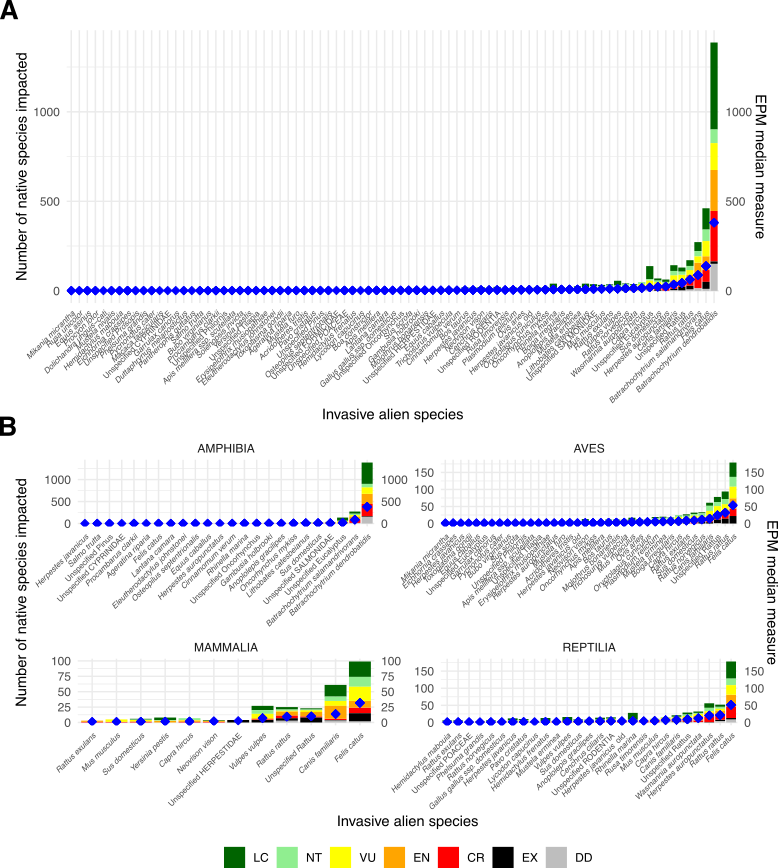

**Supplementary Figure A5. IUCN categories of native terrestrial vertebrates impacted by IAS globally and their associated EPM-A scores using low, high, and ‘NA’ impact assessments.** Only IAS with an EPM score above 1 are displayed. Panel A shows EPM median measure (blue diamonds) and the number of native species per IUCN category, based on the impacts of IAS on all native terrestrial vertebrates combined. Panel B shows EPM median measure (blue diamonds) and the number of native species per IUCN category, based on the impacts of IAS on each taxonomic group (amphibians, birds, mammals, and reptiles) separately.

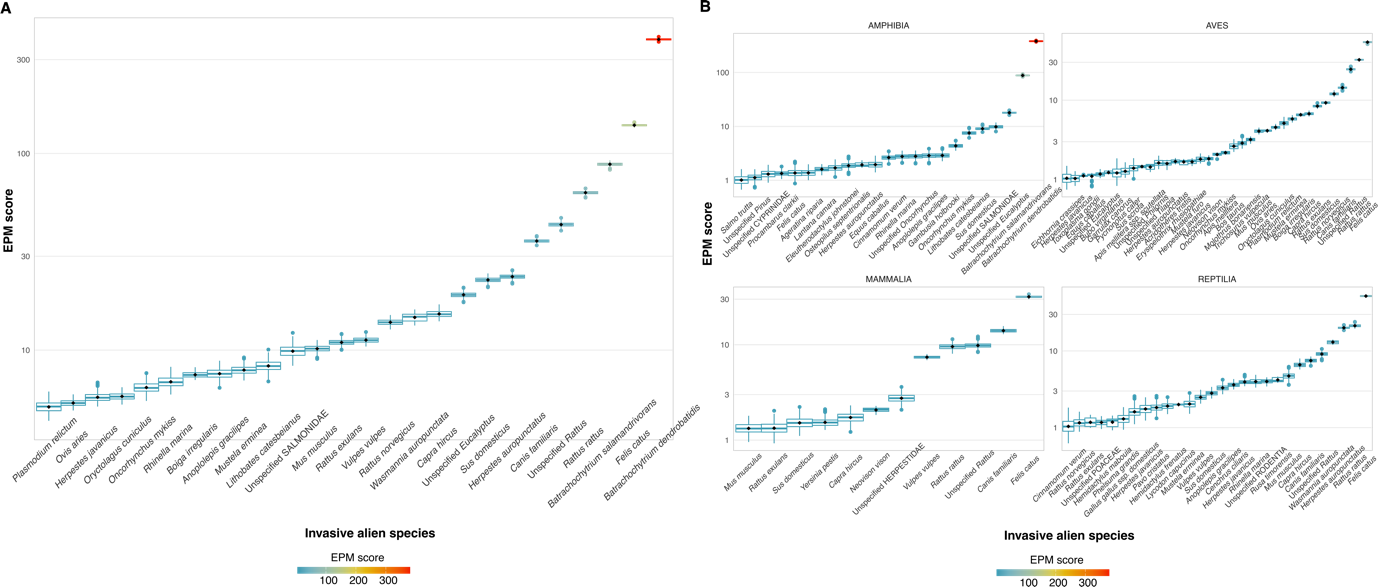

**Supplementary Figure A6.** Extinction Potential Metric (EPM-A) scores of invasive alien species (IAS) impacting native terrestrial vertebrates globally using low, high, and ‘NA’ impact assessments. Only IAS with median scores above 5 are shown in Panel A, and those with scores above 1 are shown in Panel B, for visualization purposes. Panel A shows EPM scores based on the impacts of IAS on all native terrestrial vertebrates combined. Panel B shows EPM scores based on the impacts of IAS on each taxonomic group (amphibians, birds, mammals, and reptiles) separately.

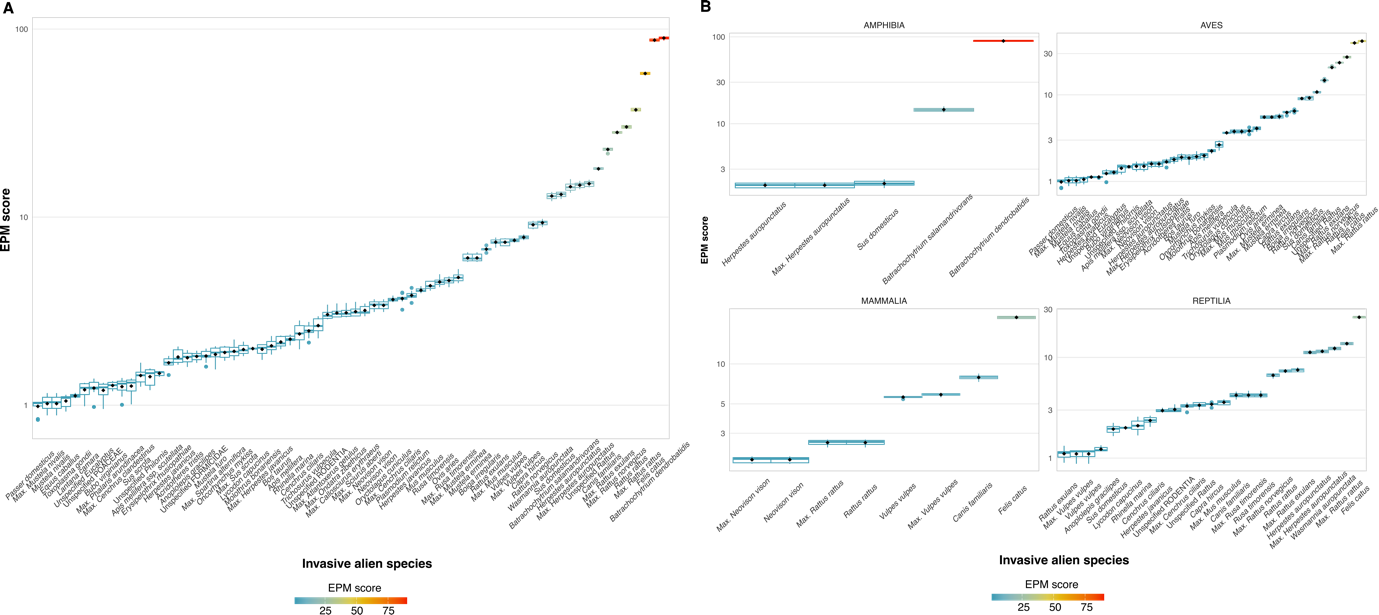
**Supplementary Figure A7. Maximum EPM-A scores of IAS impacting native terrestrial vertebrates globally using low and high impact assessments only (excluding NAs).** Only IAS with a score above 1 are displayed. Panel A shows EPM scores based on the impacts of IAS on all native terrestrial vertebrates combined. Panel B shows EPM scores based on the impacts of IAS on each taxonomic group (amphibians, birds, mammals, and reptiles) separately.

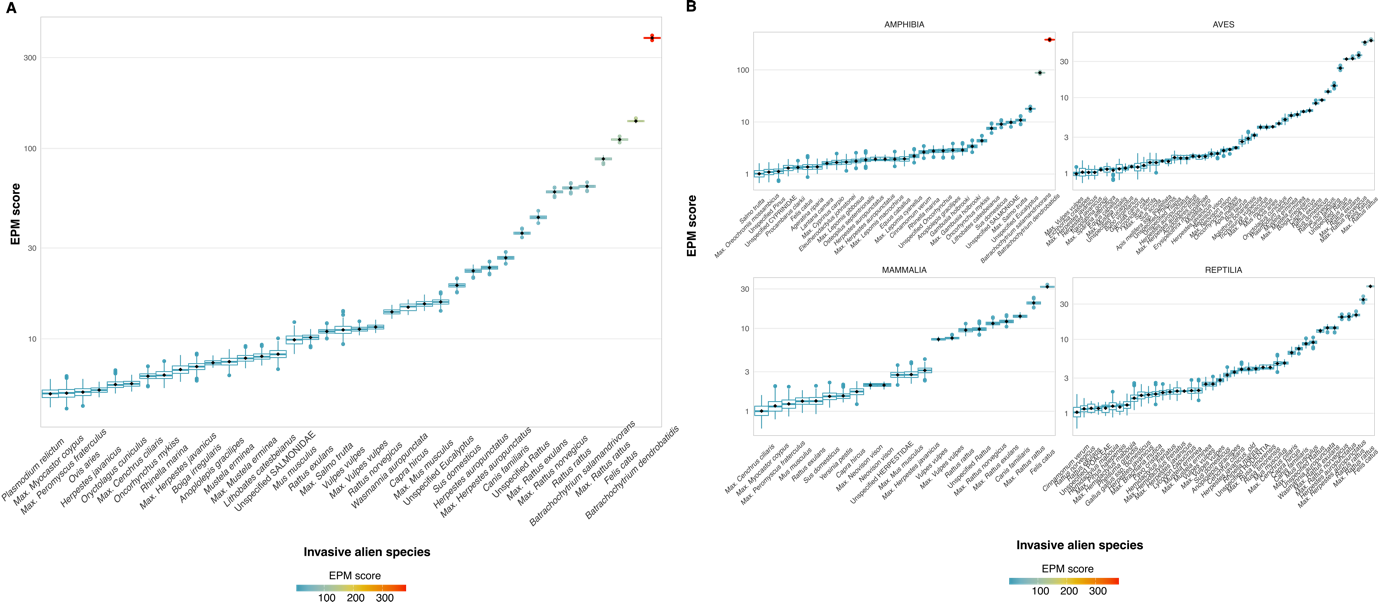

**Supplementary Figure A8.** Maximum EPM-A scores of IAS impacting native terrestrial vertebrates globally using low and high and ‘NA’ impact assessments. Only IAS with median scores above 5 are shown in Panel A, and those with scores above 1 are shown in Panel B, for visualization purposes. Panel A shows EPM scores based on the impacts of IAS on all native terrestrial vertebrates combined. Panel B shows EPM scores based on the impacts of IAS on each taxonomic group (amphibians, birds, mammals, and reptiles) separately.

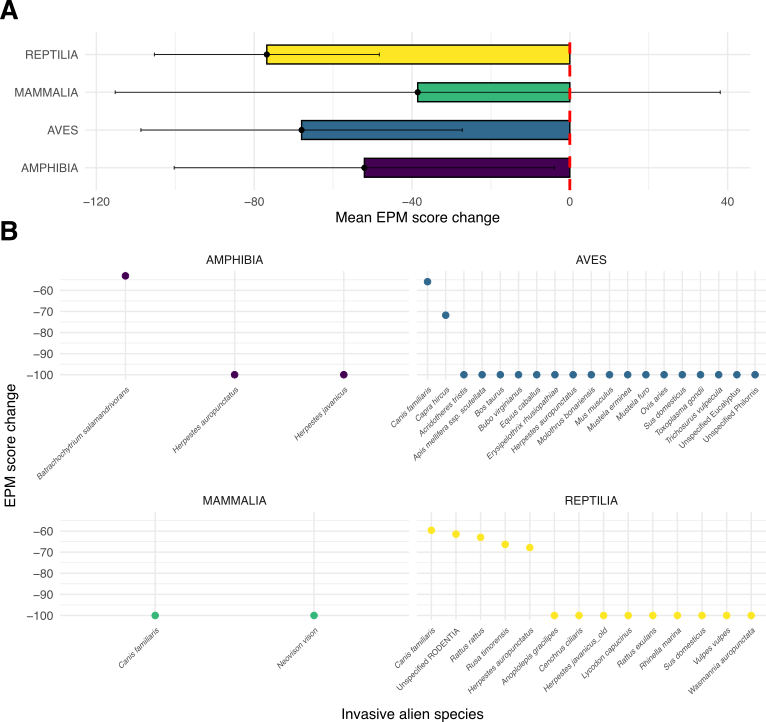

**Supplementary Figure A9. Contribution of insular endemic species to the EPM scores using low, high, and ‘NA’ impact assessments.** EPM scores reflect the impact of IAS on native terrestrial vertebrates globally, considering insular endemic species. Panel A shows the change in EPM scores (in %) when excluding insular endemic species for each of the four native taxonomic groups. Panel B highlights IAS with a difference in score of more than 50% when excluding insular endemic species.

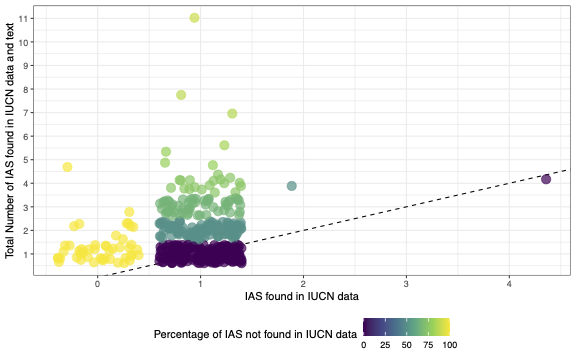

**Supplementary Figure A10. Change in total number of IAS impacting native species between tabular data and text provided by the IUCN.** Each point represents a native species and the IAS that are assessed, at least at the order level, as threatening the species. The X-axis represents the number of IAS found in the tabular data extracted from the IUCN as a text file, while the Y-axis represents the number of IAS found in both the tabular data and the justification text associated with each native species' Red List webpage. The colour indicates the percentage of IAS not found in the text, calculated using the formula: total number of IAS impacting the native species divided by the number of IAS found in the text.

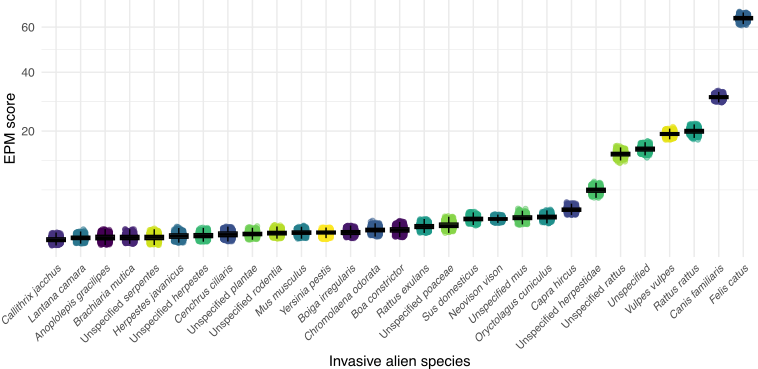

**Supplementary Figure A11. EPM-A scores of IAS impacting native mammals globally using textual and tabular data from the IUCN Red List.** EPM-A scores were computed using both textual and tabular data from the IUCN Red List, including all impacts on native species, even when the severity or scope of the impact was not assessed.

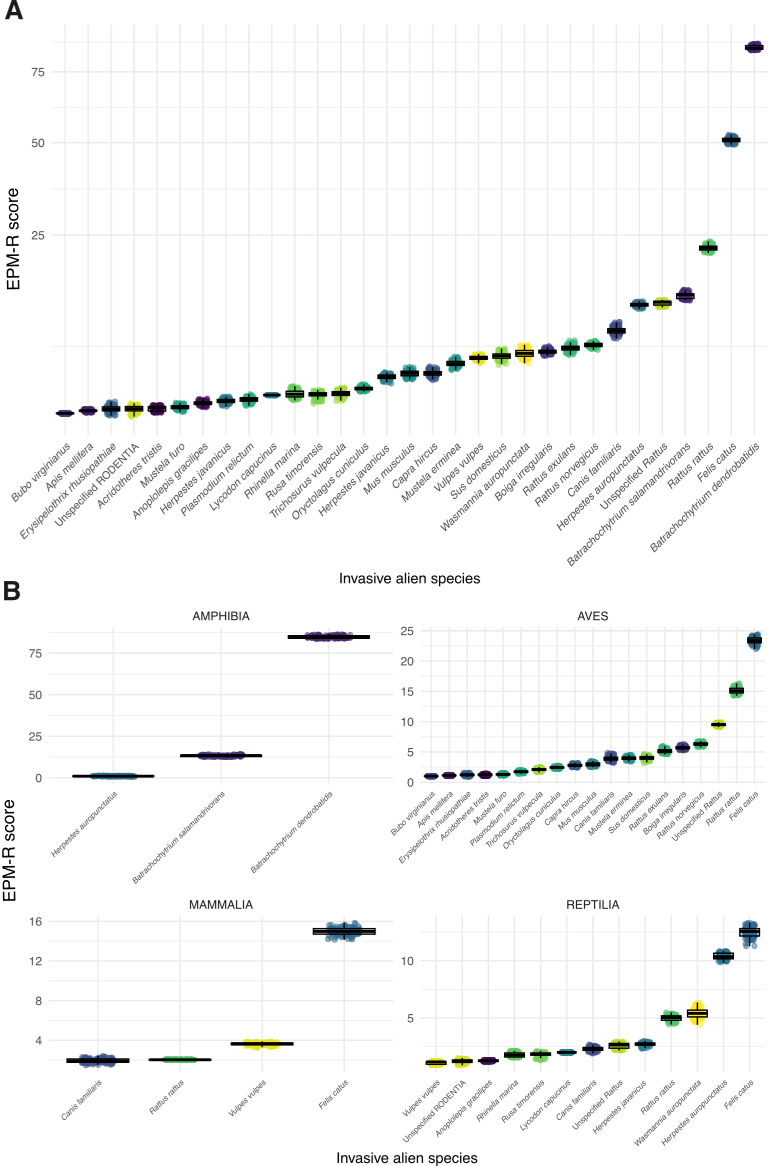

**Supplementary Figure A12. Relative Extinction Potential Metric (EPM-R) scores of IAS impacting native terrestrial vertebrates globally considering all the threats impacting native species.** Only IAS with a score above 1 are displayed. Panel A shows EPM-R scores based on the impacts of IAS on all native terrestrial vertebrates combined. Panel B shows EPM-R scores based on the impacts of IAS on each taxonomic group (amphibians, birds, mammals, and reptiles) separately.

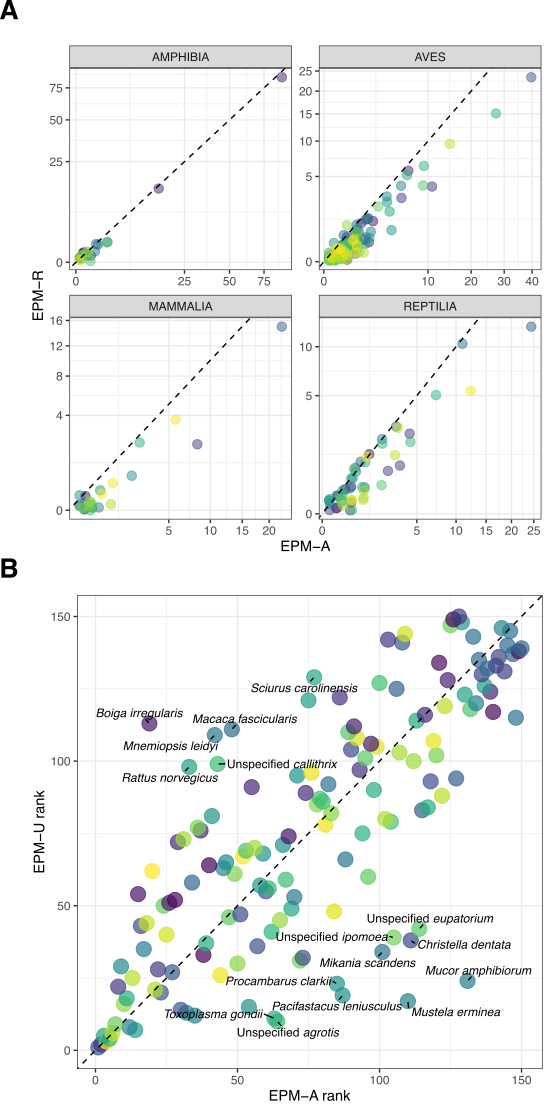

**Supplementary Figure A13. Comparison of EPM-A with EPM-R and EPM-U.** Dashed diagonal lines are 1:1 lines. Panel A, shows the EPM-A and EPM-R scores of each IAS without considering ‘NA’ impacts for each taxa. Panel B, shows the rank of IAS species according to their EPM-A and EPM-U score. The lower an IAS's rank, the higher its score, and vice versa. Species close to the diagonal have similar ranks for both approaches. IAS whose absolute difference between the two metrics is greater than 50 are displayed with their scientific name. Comparing the EPM-A and EPM-R values to examine how considering other anthropogenic changes affects the magnitude of the impact, revealing the weight of biological invasions on species extinction. By contrast, comparing EPM-A and EPM-U reveals if different IAS affect evolutionary distinct species differently.

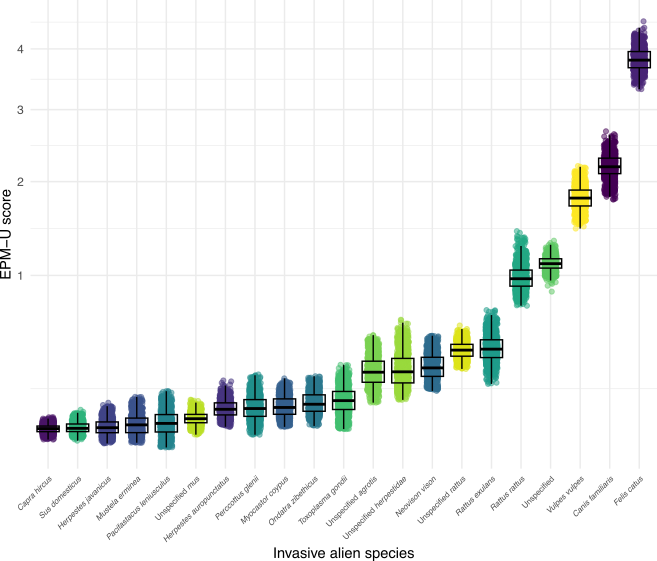

**Supplementary Figure A14. EPM-U score distributions of IAS impacting native mammals globally using low, high, and ‘NA’ impact assessments.** The points represent EPM-U values for each replicate.

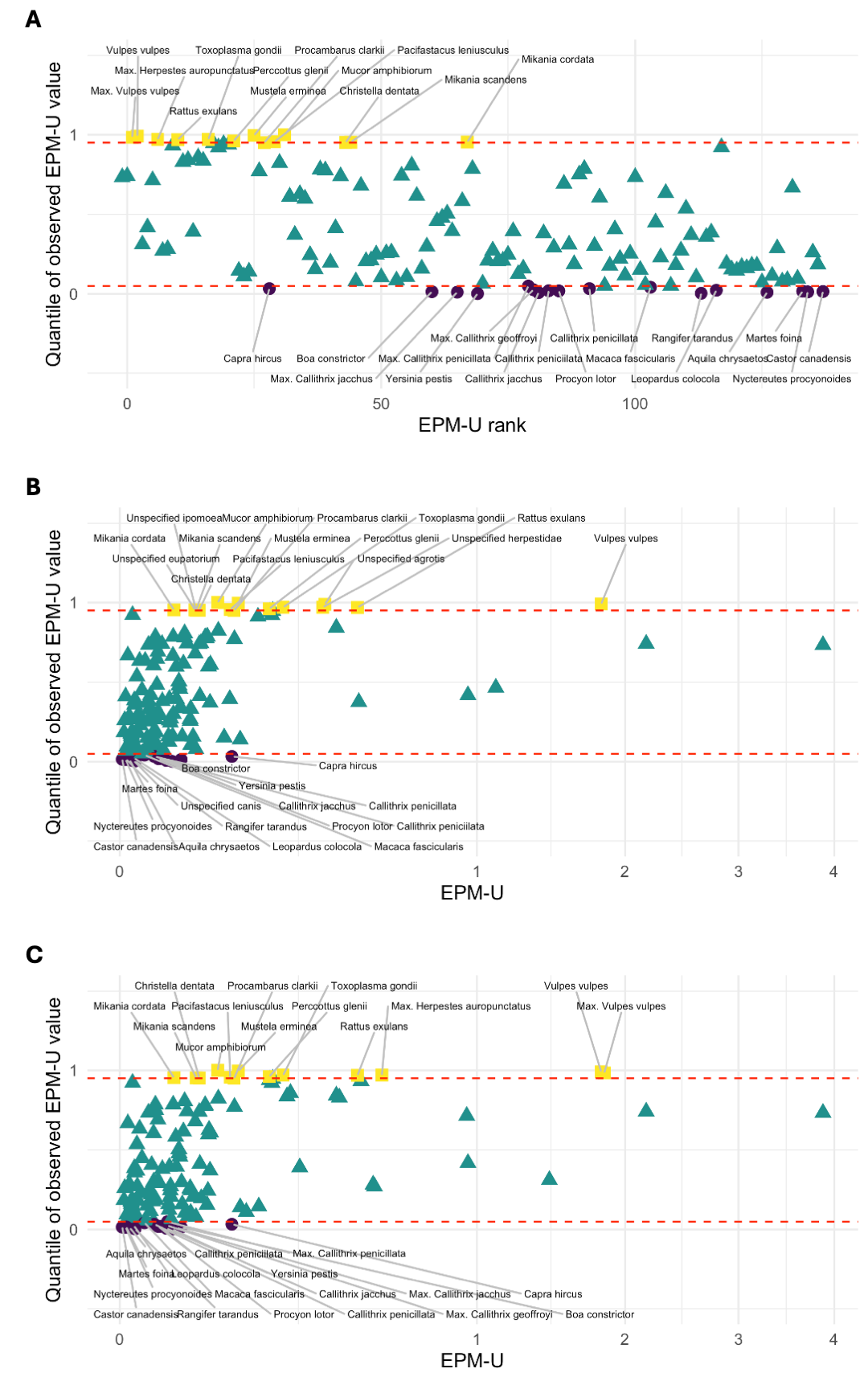

**Supplementary Figure A15. Significance of EPM-U values after randomising ED2 scores 999 times.** Panel A is similar to Fig. 6 and shows the quantiles of the maximum EPM-U scores compared to the 999 scores obtained through permutation, i.e. after accounting for unspecified species. Panels B and C plots the EPM-U scores with and without unspecified species against the EPM-U scores instead of their ranks.

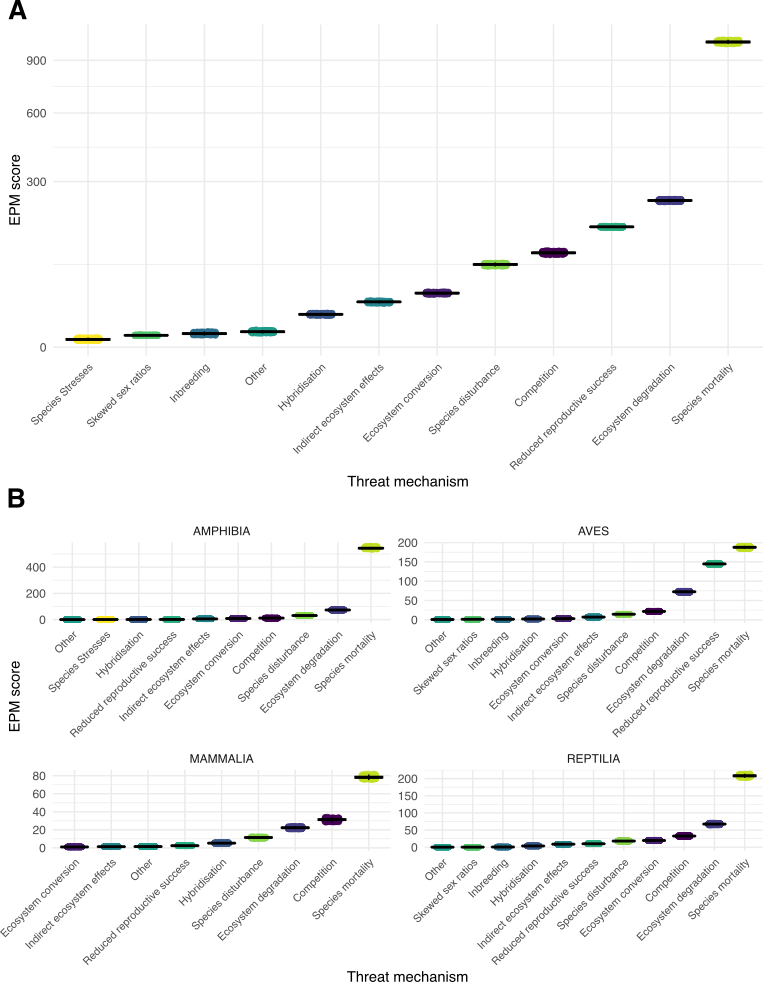

**Supplementary Figure A16. EPM scores of mechanisms impacting native terrestrial vertebrates globally using low, high, and ‘NA’ impact assessments.** Only IAS with a score above 1 are displayed. Panel A shows EPM scores based on the impacts of mechanisms on all native terrestrial vertebrates combined. Panel B shows EPM scores based on the impacts of mechanisms on each taxonomic group (amphibians, birds, mammals, and reptiles) separately.

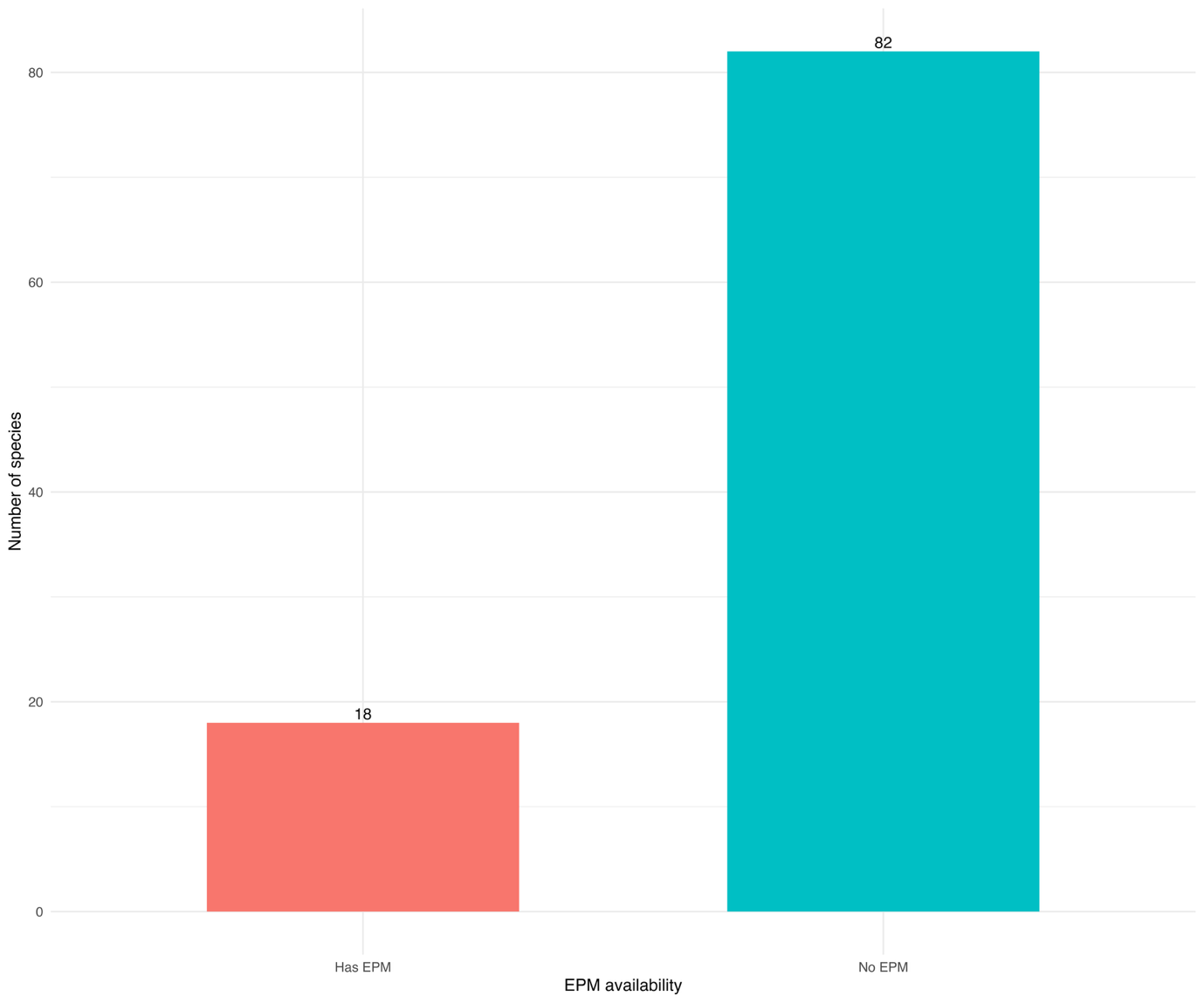

b)

a)

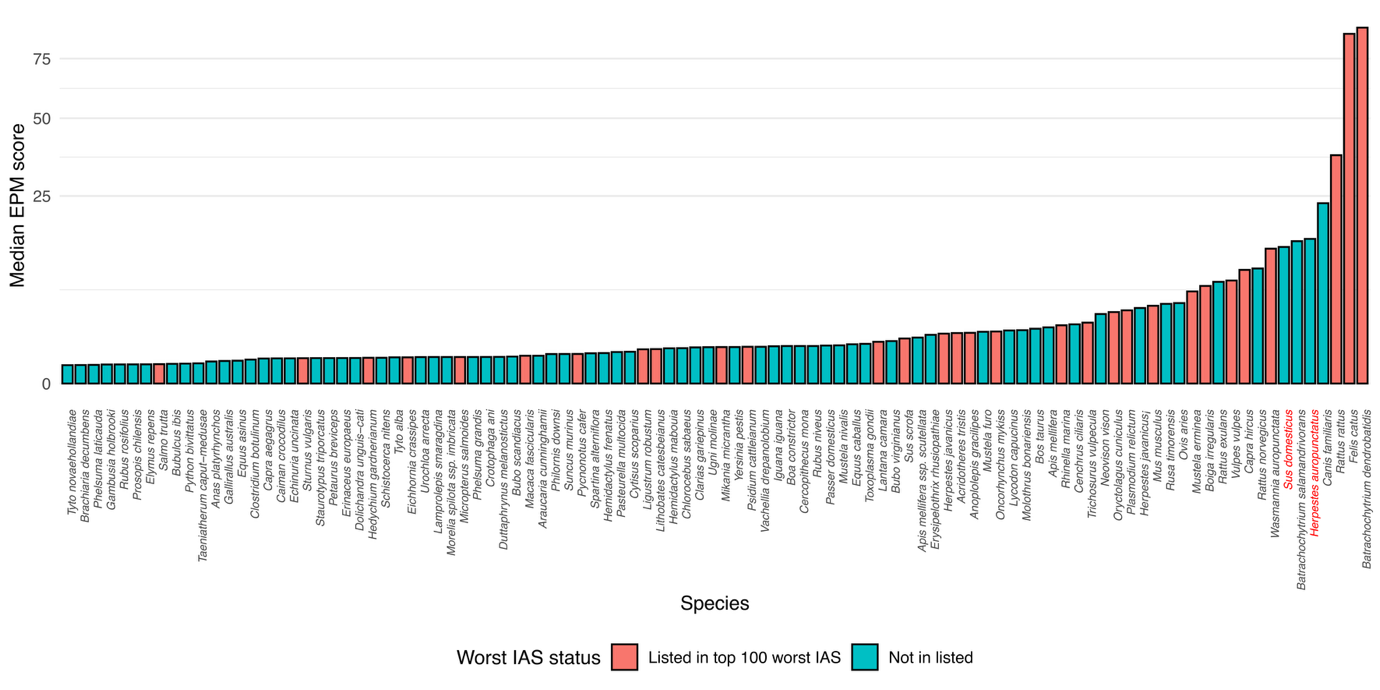

**Supplementary Figure A17.** EPM values for species on the 100 of the worst invasive species list. a) Number of species on the 100 list with an EPM value (i.e. listed as a threat in the IUCN Red List data). b) EPM values of the species on the 100 list. *Sus domesticus* was considered to be on the list as a subspecies of *Sus scrofa*.

**APPENDIX B: Modelling the empirical distributions of EPM-A values**

**Methods**

For each set of EPM-A values (each taxonomic group and considering or not NA impact magnitudes), we modelled the empirical distribution of impact values by fitting four standard continuous families—power law, lognormal, Weibull, and exponential— to examine the underlying processes behind impacts, and to examine if impact distributions vary across taxonomic groups (suggesting different processes at play). These four distributions span the full spectrum of tail behaviour, from very heavy tails (power law) to very light (exponential). A light tail would suggest large impacts to be rare, whereas a heavy tail would indicate high impacts to be relatively common.

The distributions were fitted to the whole dataset, and to the upper tail only (i.e. each distribution was fitted twice). The tail was analysed separately because heavy‑tailed behaviour, if present, typically holds only beyond a threshold (Clauset et al. 2009). We identified this threshold (xmin) using the Clauset–Shalizi–Newman approach, which applies xmin to minimise the Kolmogorov–Smirnov distance for a power‑law fit; all tail models were then fit by maximum likelihood on the left‑truncated data (x ≥ xmin), and the whole‑range models were fit by maximum likelihood on all observations. We compared models for each taxonomic group using the corrected Akaike Information Criterion (AICc).

**Results**

The upper tails of the distributions of EPM-A values were overall better described by power laws (the widest tail of the fitted distributions) than by lognormal, Weibull or exponential distributions, with the lightest-tail exponential consistently having the highest AICc values for all taxonomic groups (Supp. Table B1), indicating that large impacts are not uncommon. The tail fraction (the number of IAS with an impact above xmin, divided by the total number of IAS) tended to be larger for mammals and reptiles than for amphibians and birds, although that depended partly on the version of EPM-A used (i.e. considering or excluding unspecified species or data deficient impact magnitudes). This indicates that a larger proportion of IAS are included in the tail and have very large impacts on native species for reptiles and mammals.

A lognormal distribution was the most common fit for the whole distribution (i.e. without considering cutoff values), indicating most IAS cause small to moderate impacts, spread over a wide range.

**Supplementary Table B1.** AIC values after fitting Power Law (pl), Lognormal (ln), Weibull (wb) and Exponential (exp) distributions to the EPM-A values, considering raw data, unspecified species, and including or excluding data deficiencies for impact magnitude (excluding: no_DD). “whole_model” columns are the results after fitting the distributions for all IAS, and “tail_model” columns are the results after fitting the distributions only for the most impactful species, after determining the cutoff value x_min, as described above (tail_n is the number of IAS included after applying the xmin threshold, and tail_fraction is tail_n divided by the total number of IAS).

| EPM-A version | Taxonomic group | best_whole_model | AICc_pl_whole | AICc_ln_whole | AICc_wb_whole | AICc_exp_whole | xmin | tail_n | tail_fraction | best_tail_model | AICc_pl_tail | AICc_ln_tail | AICc_wb_tail | AICc_exp_tail |
| --- | --- | --- | --- | --- | --- | --- | --- | --- | --- | --- | --- | --- | --- | --- |
| EPM-A | AMPHIBIA | **Lognormal** | 252.201 | **217.450** | 265.527 | 532.650 | 0.932 | 25 | 0.263 | **Power law** | **124.922** | 127.338 | 130.465 | 204.356 |
| EPM-A | AVES | **Lognormal** | 364.223 | **300.301** | 351.820 | 505.560 | 0.936 | 44 | 0.259 | **Power law** | **179.311** | 181.541 | 182.763 | 221.897 |
| EPM-A | MAMMALIA | **Lognormal** | 116.188 | **103.650** | 135.982 | 213.467 | 0.307 | 47 | 0.534 | **Power law** | **79.214** | 81.424 | 85.870 | 155.750 |
| EPM-A | REPTILIA | **Lognormal** | 261.962 | **237.693** | 264.990 | 375.825 | 0.460 | 61 | 0.508 | **Power law** | **185.486** | 187.243 | 188.250 | 251.265 |
| EPM-A NO_DD | AMPHIBIA | **Power law** | **45.533** | 52.896 | 61.1519 | 111.884 | 0.264 | 12 | 0.6 | **Power law** | **45.774** | 48.903 | 49.773 | 79.7438 |
| EPM-A NO_DD | AVES | **Lognormal** | 317.993 | **262.872** | 300.282 | 409.417 | 0.391 | 74 | 0.514 | **Power law** | **208.533** | 209.816 | 210.564 | 280.743 |
| EPM-A NO_DD | MAMMALIA | **Power law** | **39.736** | 50.352 | 60.0427 | 85.0282 | 0.143 | 19 | 0.730 | **Power law** | **41.413** | 43.976 | 44.412 | 72.028 |
| EPM-A NO_DD | REPTILIA | **Lognormal** | 184.197 | **151.629** | 156.0286 | 176.794 | 0.395 | 35 | 0.686 | **Weibull** | 118.627 | 118.684 | **118.463** | 136.587 |
| EPM-A Unspecified spp | AMPHIBIA | **Lognormal** | 313.269 | **279.791** | 335.4712 | 618.687 | 0.365 | 72 | 0.615 | **Power law** | **209.838** | 211.984 | 217.558 | 443.439 |
| EPM-A Unspecified spp | AVES | **Lognormal** | 483.703 | **405.211** | 473.804 | 710.873 | 0.479 | 95 | 0.466 | **Power law** | **305.965** | 308.032 | 312.024 | 446.528 |
| EPM-A Unspecified spp | MAMMALIA | **Lognormal** | 222.351 | **190.780** | 224.9845 | 324.737 | 0.383 | 53 | 0.477 | **Power law** | **157.470** | 159.441 | 160.517 | 215.668 |
| EPM-A Unspecified spp | REPTILIA | **Lognormal** | 401.151 | **359.375** | 391.520 | 536.098 | 0.468 | 81 | 0.54 | **Power law** | **280.760** | 282.117 | 282.941 | 364.553 |
| EPM-A Unspecified spp NO_DD | AMPHIBIA | **Power law** | **55.6925** | 61.823 | 72.7568 | 133.133 | 0.264 | 16 | 0.615 | **Power law** | **52.968** | 56.280 | 57.152 | 97.332 |
| EPM-A Unspecified spp NO_DD | AVES | **Lognormal** | 408.583 | **347.023** | 398.528 | 569.384 | 0.448 | 81 | 0.463 | **Power law** | **272.201** | 273.562 | 274.537 | 361.138 |
| EPM-A Unspecified spp NO_DD | MAMMALIA | **Power law** | **54.102** | 67.707 | 77.610 | 105.708 | 0.142 | 25 | 0.758 | **Power law** | **56.927** | 59.194 | 59.477 | 90.957 |
| EPM-A Unspecified spp NO_DD | REPTILIA | **Lognormal** | 263.586 | **221.508** | 224.978 | 252.692 | 0.395 | 47 | 0.701 | **Weibull** | 182.304 | 180.305 | **179.395** | 197.897 |

**APPENDIX C: Comparing past and future impacts of different magnitudes**

**Methods**

EPM-A incorporates information on past and projected future extinctions, to combine different levels of impact magnitude in a single metric. We explored the relationship between past extinctions and future extinctions (i.e. current impacts on native populations) over 50 years, further separating native species heavily and moderately impacted by IAS. To do so, for each IAS, we computed EPM-A including native species in the EX and EW Red List categories only (i.e. the number of native species that went extinct because of the IAS; EPM-A-Past hereafter), EPM-A for native species in the CR category (i.e. native species with a high probability of extinction; EPM-A-High hereafter) and EPM-A for native species in the EN, VU, NT and LC categories (i.e. native species with a lower probability of extinction; EPM-A-Low hereafter). We then computed the difference between each pair of these three EPM-A values for each IAS (EPM-A-High – EPM-A-Past, EPM-A-Low – EPM-A-Past, EPM-A-Low – EPM-A-High), using median values across replicates. Positive values indicate that future impacts will be higher than past ones in terms of species extinction. This was done for all native taxonomic groups together, and for each group separately.

We then explored if species with high impacts in the past also had high impacts in terms of probabilities of extinction, and if these relationships are linear. To do so, for each pairwise combination the three metrics, we first removed IAS with EPM-A-Past, EPM-A-High and EPM-A-Low values of 0. This is a conservative approach that assumes that a values of 0 is likely to result from a lack of assessment, or that recently introduced species will likely not have had the time to lead species to extinction (or to the brink of extinction). Future, more in-depth analyses would need to examine history of introduction and reporting biases for all species. We then computed the regression between each pairwise combination after log-transforming these values. A coefficient of 1 would indicate the relationship is linear; a coefficient <1 indicates saturation (species which had major impacts in the past will affect fewer species in the future, either because they have since been managed or because they have already impacted species they were going to impact); and a coefficient >1 means that there will be an acceleration (species which had major impacts in the past will have disproportionately higher impacts in the future). To assess if the coefficients were statistically different from 1, we computed the following linear models and examined if the coefficients were significantly different from 0:

$\log\left( \text{EPM-A-High} \right)-\log\left( \text{EPM-A-Past} \right)\sim\log\left( \text{EPM-A-Past} \right)$	Eq.1

$\log\left( \text{EPM-A-Low} \right)-\log\left( \text{EPM-A-Past} \right)\sim\log\left( \text{EPM-A-Past} \right)$	Eq.2

$\log\left( \text{EPM-A-Low} \right)-\log\left( \text{EPM-A-High} \right)\sim\log\left( \text{EPM-A-High} \right)$	Eq.3

Due to data limitation after removing 0 values, this was only done for all native taxonomic groups combined.

**Results**

Differences between the three types of EPM-A values (EPM-A-High – EPM-A-Past, EPM-A-Low – EPM-A-Past, EPM-A-Low – EPM-A-High) were skewed towards positive values for all taxonomic groups, indicating that future impacts will be higher than past ones (i.e. EPM-A-Low > EPM-A-High > EPM-A-Past overall), although most differences were small, between [0,1]. This was true when considering unspecified species as separate species and for maximum EPM-A values, and when including or excluding threats whose impact magnitude was classified as “not available” (NA).

Regression coefficients from Eqs 1-3 were not significantly different from 0 for the slopes, indicating that the relationships between EPM-A-Past, EPM-A-High and EPM-A-Low were linear (Supp. Fig. C2, Supp. Table C1).

**Supplementary Table C1.** Coefficients with standard errors and p-values for linear models described in Eqs 1-3. Model 1: unspecified species considered as separate species; Model 2: unspecified species considered as separate species and threats whose impact magnitude was classified as “not available” (NA) excluded; Model 3: maximum EPM-A values; Model 4: maximum EPM-A values and threats whose impact magnitude was classified as “not available” (NA) excluded.

| Model # |  | Past-High | | | Past-Low | | | High-Low | | |
| --- | --- | --- | --- | --- | --- | --- | --- | --- | --- | --- |
|  |  | Coef | SE | p-value | Coef | SE | p-value | Coef | SE | p-value |
| 1 | Intercept | 0.396 | 0.274 | 0.163 | -0.114 | 0.299 | 0.707 | **-0.451** | **0.147** | **0.003** |
|  | Predictor | -0.082 | 0.189 | 0.668 | 0.191 | 0.211 | 0.374 | 0.175 | 0.112 | 0.121 |
| 2 | Intercept | 0.417 | 0.254 | 0.112 | -0.028 | 0.255 | 0.912 | **-0.307** | **0.129** | **0.020** |
|  | Predictor | -0.130 | 0.148 | 0.388 | 0.133 | 0.155 | 0.398 | 0.127 | 0.094 | 0.150 |
| 3 | Intercept | 0.040 | 0.457 | 0.931 | -0.483 | 0.262 | 0.082 | -0.063 | 0.204 | 0.761 |
|  | Predictor | -0.067 | 0.302 | 0.827 | 0.305 | 0.195 | 0.135 | -0.096 | 0.161 | 0.557 |
| 4 | Intercept | 0.273 | 0.413 | 0.517 | -0.324 | 0.245 | 0.199 | -0.084 | 0.174 | 0.631 |
|  | Predictor | -0.232 | 0.252 | 0.369 | 0.246 | 0.165 | 0.149 | -0.097 | 0.135 | 0.474 |

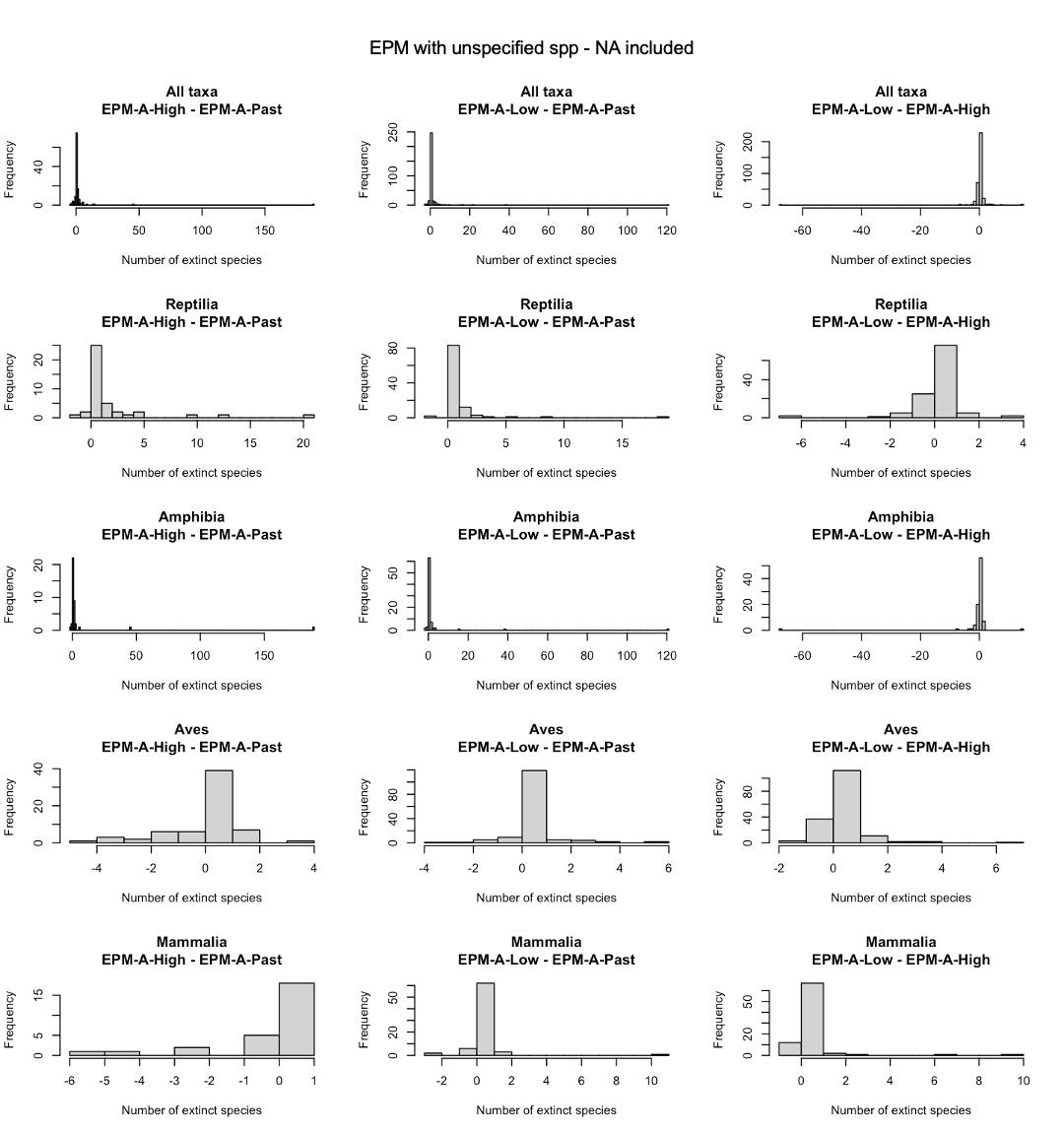

**Supplementary Figure C1.** Histograms of the differences of median EPM-A values computed for EX/EW (past extinctions, EPM-A-Past), CR (high impacts, EPM-A-High) and EN/VU/NT/LC (low impacts, EPM-A-Low) categories, considering unspecified species as separate species. Each column is a different pairwise association. Each row is a different taxonomic group. Positive values indicate that the sum of high impacts is larger than the sum of past impacts (column 1), or that the sum of low impacts is larger than the sum of past (column 2) or high (column 3) impacts.

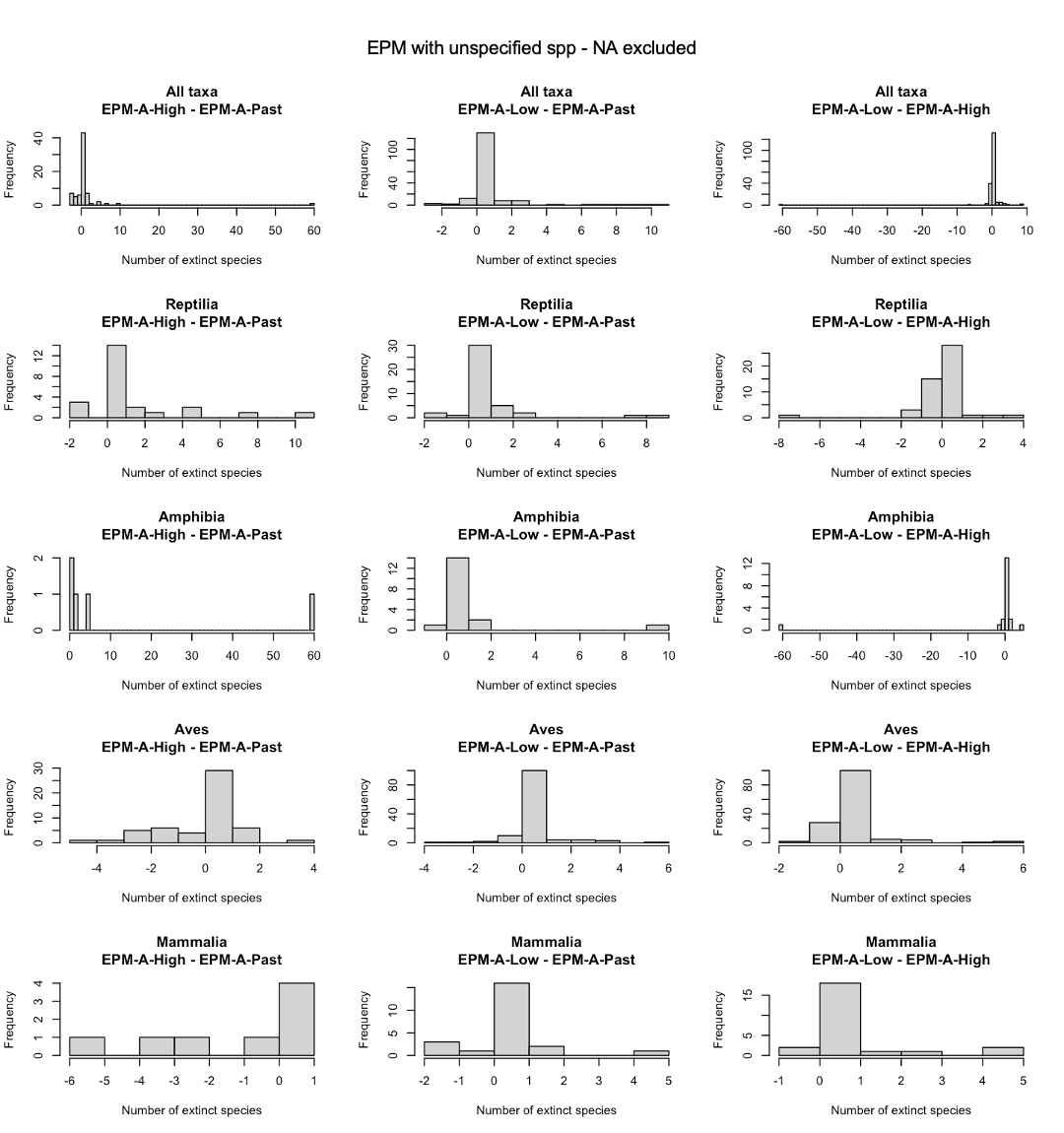

**Supplementary Figure C2.** Histograms of the differences of median EPM-A values computed for EX/EW (past extinctions, EPM-A-Past), CR (high impacts, EPM-A-High) and EN/VU/NT/LC (low impacts, EPM-A-Low) categories, considering unspecified species as separate species and after excluding threats whose impact magnitude was classified as “not available” (NA). Each column is a different pairwise association. Each row is a different taxonomic group. Positive values indicate that the sum of high impacts is larger than the sum of past impacts (column 1), or that the sum of low impacts is larger than the sum of past (column 2) or high (column 3) impacts.

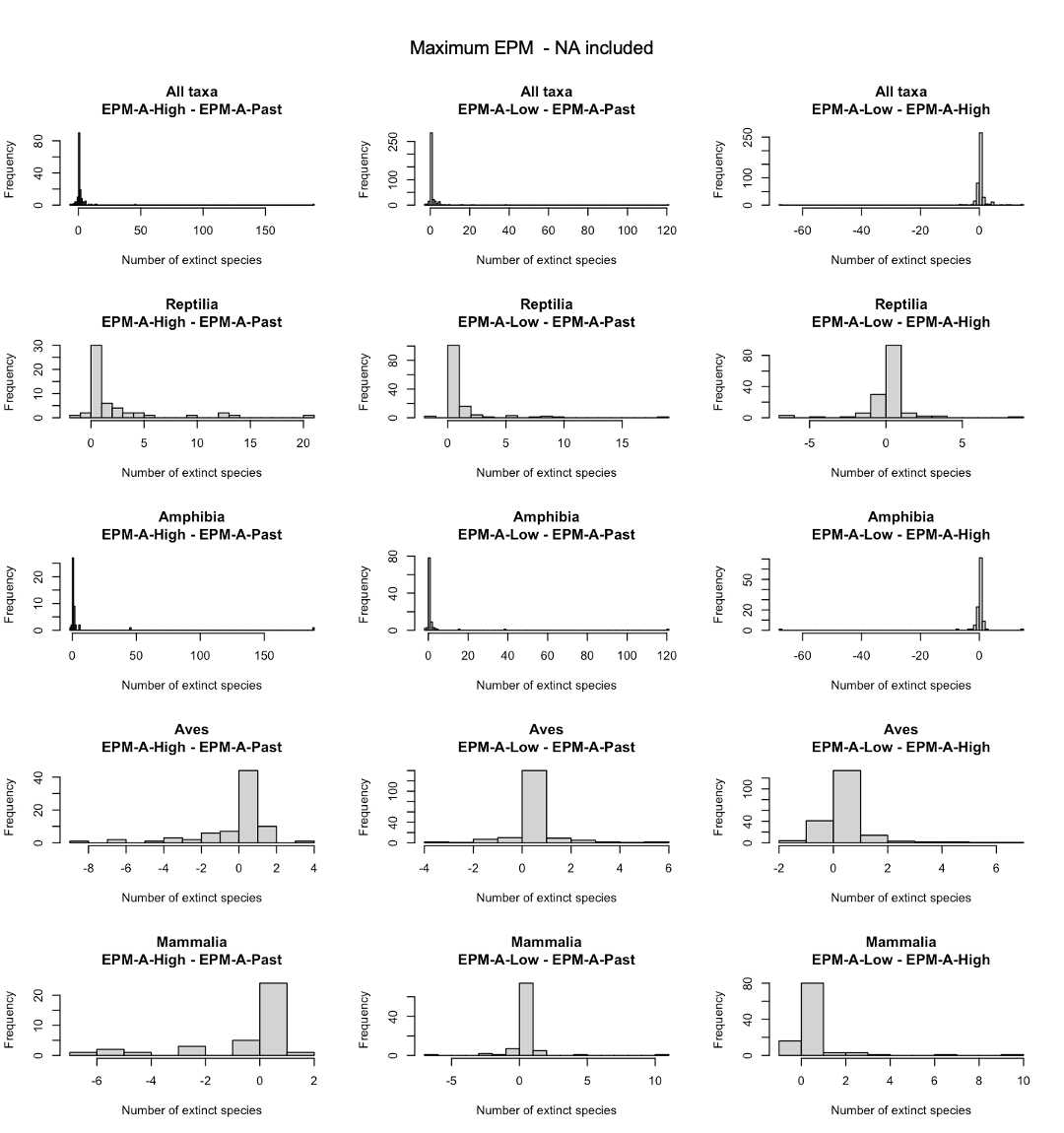

**Supplementary Figure C3.** Histograms of the differences of median EPM-A values computed for EX/EW (past extinctions, EPM-A-Past), CR (high impacts, EPM-A-High) and EN/VU/NT/LC (low impacts, EPM-A-Low) categories, for maximum EPM-A values. Each column is a different pairwise association. Each row is a different taxonomic group. Positive values indicate that the sum of high impacts is larger than the sum of past impacts (column 1), or that the sum of low impacts is larger than the sum of past (column 2) or high (column 3) impacts.

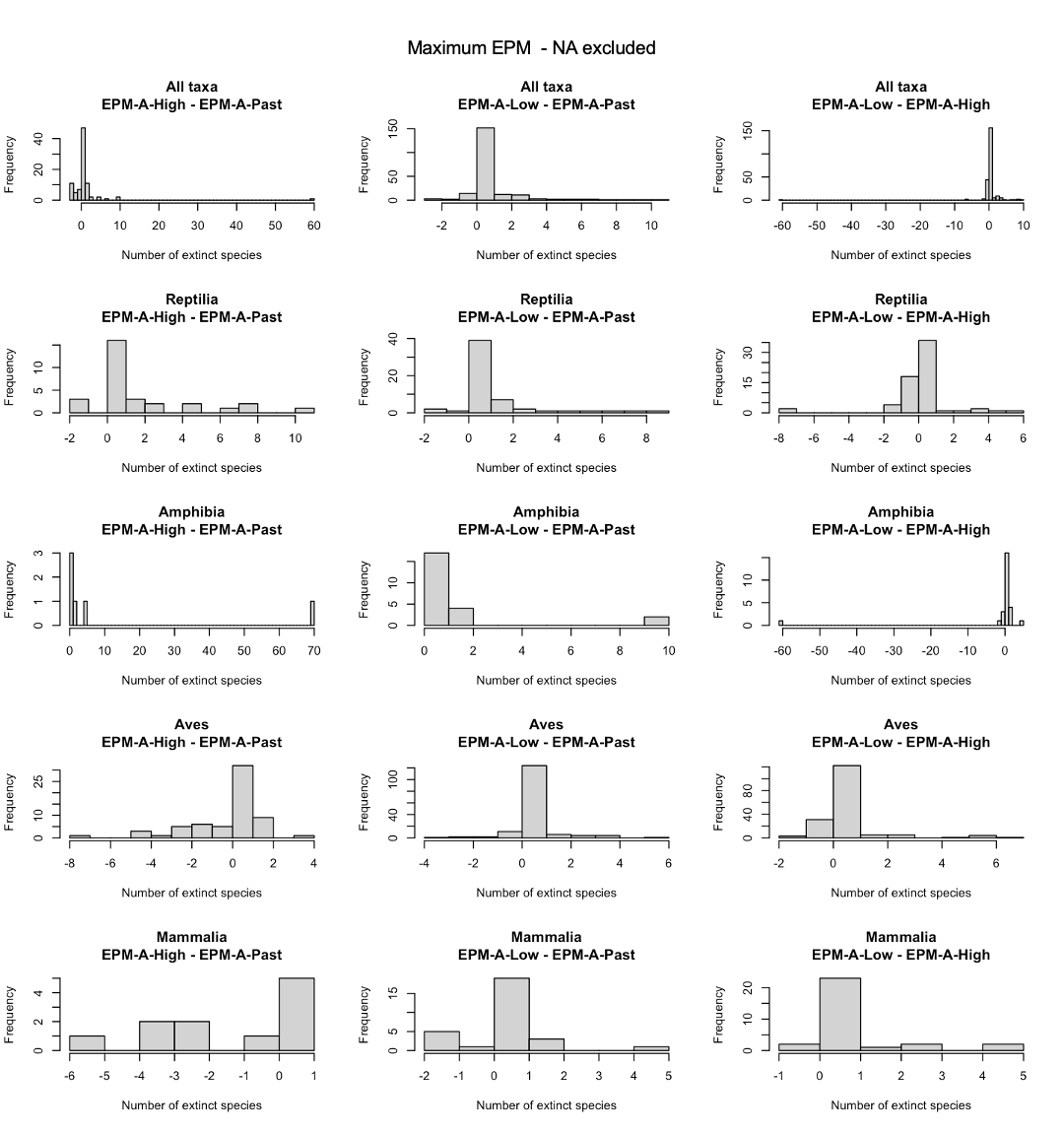

**Supplementary Figure C4.** Histograms of the differences of median EPM-A values computed for EX/EW (past extinctions, EPM-A-Past), CR (high impacts, EPM-A-High) and EN/VU/NT/LC (low impacts, EPM-A-Low) categories, for maximum EPM-A values and after excluding threats whose impact magnitude was classified as “not available” (NA). Each column is a different pairwise association. Each row is a different taxonomic group. Positive values indicate that the sum of high impacts is larger than the sum of past impacts (column 1), or that the sum of low impacts is larger than the sum of past (column 2) or high (column 3) impacts.

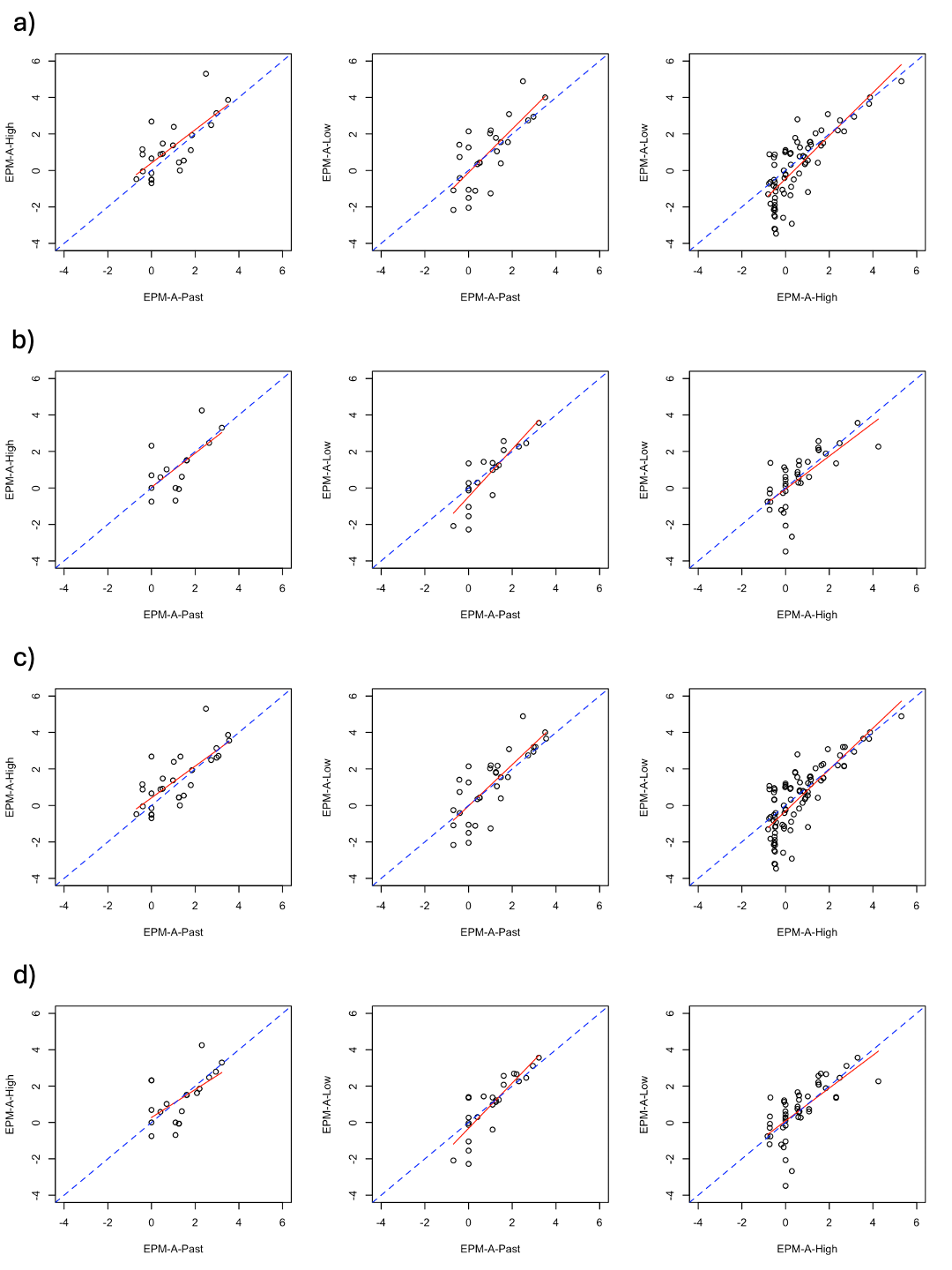

**Supplementary Figure C5.** Relationship between EPM-A values computed for EX/EW (past extinctions, EPM-A-Past), CR (high impacts, EPM-A-High) and EN/VU/NT/LC (low impacts, EPM-A-Low) categories. a) Unspecified species considered as separate species; b) Unspecified species considered as separate species and threats whose impact magnitude was classified as “not available” (NA) excluded; c) Maximum EPM-A values; d) Maximum EPM-A values and threats whose impact magnitude was classified as “not available” (NA) excluded. Values on the x and y-axes have been log-transformed. The dashed, blue lines represent the theoretical 1:1 relationship. The red, continuous lines represents the fitted relationships.
